# Ancient inversion polymorphisms are locally adaptive in a widespread butterfly species

**DOI:** 10.1101/2024.09.20.614156

**Authors:** Fernando Seixas, Sarah Dendy, Shuzhe Guan, Neil Rosser, Nick Grishin, Neil Davies, Lawrence E. Gilbert, W. Owen McMillan, James Mallet

## Abstract

Wide-ranging species are subject to varying biotic and abiotic selection pressures across their distribution. While local adaptation does not manifest in obvious morphological changes, population genomic studies can reveal cryptic diversity and provide insights into local adaptive processes. In this study, we investigated the biogeographic history and genomic diversity across the range of the zebra longwing butterfly *Heliconius charithonia,* a species with a widespread distribution in the Neotropics, but which is phenotypically homogenous across its range. We examined whole genome sequence data from 55 individuals from the eight described subspecies. We infer that there were at least two distinct colonization events of the Caribbean islands from the mainland. The second colonization wave occurred relatively recently, accounting for the genetic homogeneity observed across the species’ range. Despite low divergence across most of the genome, two large non-recombining genomic regions showed deeply divergent haplotypes that correspond to chromosomal inversions. Phylogenetic analyses indicate ancient origins of these inversion polymorphisms, and there is no evidence that they were introgressed from another extant lineage of *Heliconius*. These ancient polymorphisms are likely maintained by heterogeneous selection across the landscape, with the inversion on chromosome 19 likely playing a role in local adaptation to cold and desiccation. Our findings underscore the importance of genomic analysis in uncovering hidden diversity and adaptation in phenotypically homogenous species and highlight the significant role of chromosomal inversions in driving local adaptation.

## Introduction

Species with extensive distributions are exposed to diverse selective pressures across their ranges. This variation arises from heterogeneity of biotic factors, such as competition and predation, as well as of abiotic conditions like climate, geography, and availability of resources. Species must adapt to a multitude of environmental challenges and opportunities, leading to a complex and heterogeneous selective landscape that can drive local adaptation and influence evolutionary trajectories.

Adaptation in the face of gene flow can be achieved provided divergent selection is strong enough to prevent the loss of advantageous alleles and maintain a migration-selection equilibrium; in contrast to traditional views in animal evolutionary biology ^1^, gene flow has a relatively weak effect on local divergence across species ranges ^2–4^. However, local adaptation can be challenging when it involves alleles at multiple loci, since gene flow can break up adaptive combinations ^5^. When adaptation is very local compared to dispersal range, genetic architectures that reduce recombination between loci involved in adaptive traits are likely to evolve ^6,7^. Chromosomal inversions are important modifiers of the recombination landscape. When heterozygous, inversions usually reduce recombination between standard and inverted haplotypes and can maintain linkage disequilibrium among adaptive allele and facilitate adaptive divergence in the face of gene flow ^5^.

Traditional approaches to studying adaptation often focus on directly observable, often morphological, traits. While such studies can be powerful and can lead to clues about the genetic basis of an adaptation, they can also introduce bias in understanding the relative role of different adaptive processes and underlying genomic architecture. In contrast, bottom-up genomic approaches, which examine genetic variation without relying on phenotypic assumptions, offer several advantages. Being agnostic to phenotypes an unbiased approach avoids *a priori* hypotheses about which groups of populations or traits are under selection; this can be particularly helpful in organisms in which the natural history is not well studied and in species with cryptic genetic variation. Additionally, they provide a comprehensive and unbiased means to identify adaptive processes and their genetic underpinnings ^8^. For instance, while many studies have linked adaptive phenotypes to structural variants like large inversions, the ease of detecting these inversions – due to their ability to reduce recombination and promote divergence – raises the question of whether their importance in adaptation results from detection bias ^9^.

The zebra longwing butterfly, *Heliconius charithonia*, is widely distributed across the Caribbean and Gulf of Mexico (Figure 1). Its range extends from South America to southern United States, and to the Greater Antilles. This species is unique, in that it is the only *Heliconius* species found on the major Caribbean islands, which suggests dispersal abilities lacked by other *Heliconius* species. Unlike other *Heliconius*, which are iconic for the geographic diversity of wing color pattern, *H. charithonia* varies little across its extensive geographic range. Although there are currently eight recognized subspecies ^10,11^, these can be recognized only based on minor differences in color pattern. Genetic diversity and spatial genetic structure in *H. charithonia* remain poorly described. Genetic variation across the majority of the species range has been studied only once, using low-resolution data from two mitochondrial genes, restriction-fragment-length polymorphisms, and allozyme data ^12^. Relationships between populations was often poorly resolved due to low levels of divergence (0.4%) at mtDNA, suggesting a recent and rapid colonization of the Caribbean. The Jamaican subspecies was an exception in that it appeared to be basal to the group and very distinct from other populations (2.4% at mtDNA), likely a result of an earlier colonization.

**Figure 1.**
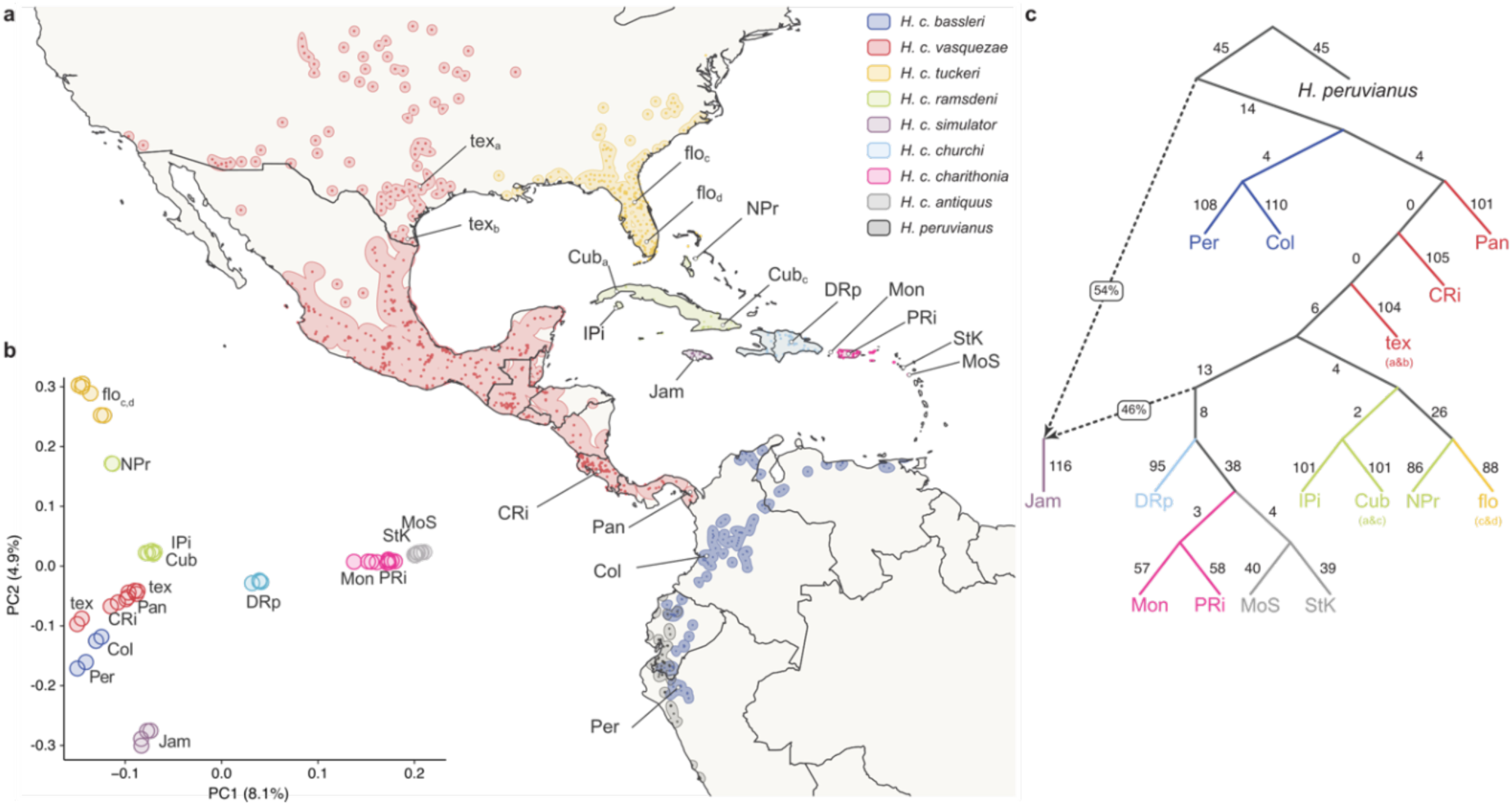
- Sampling and population structure of *H. charithonia*. **(a)** *H. charithonia* species distribution and sampling locations (coordinates and sampling location code names are provided in Supplementary Table 1). Subspecies’ ranges are depicted in different colors and were inferred based on historical and current records (https://heliconius-maps.github.io/index.html). **(b)** Principal component analysis (PCA) based on 12,342 autosomal SNPs. **(c)** An admixture-graph model of *H. charithonia* suggests two colonization waves of the Caribbean. Dashed edges indicate admixture events, with ancestry proportions as percentages within boxes. Solid edges and corresponding numbers indicate drift.

Here we investigate the biogeographic history of *H. charithonia* and characterize genetic diversity across the geographic range of the species using whole genome sequence data. We first examine population structure and phylogenetic relationships to reconstruct the colonization history of the Caribbean (the number of colonization waves, their origin and migration routes). We then explore patterns of genetic diversity and recombination across the genome and identify putative inversion polymorphisms. Finally, we investigate the evolutionary history of these inversions and the role of selection in the maintenance of inversion polymorphisms, namely associations between inversion haplotypes and environmental conditions across the species range.

## Results

### Population structure and biogeographic history of H. charithonia

We analyzed whole-genome re-sequencing data from 54 *Heliconius charithonia*, sampled from 17 locations spanning the species’ range and including representatives of all eight subspecies (Figure 1a; Supplementary Table 1). Individuals group according to their putative subspecies and geographic location, based on principal component analysis (PCA) of single nucleotide polymorphisms (SNPs, Figure 1b). The first two first principal components explain only 13% of the total variation due to an overall lack of strong population structure (Figure 1b). Broadly, the first component distinguishes between continental and island populations and the second component between populations north of Cuba (i.e. New Providence and Florida) and east of Cuba (Puerto Rico, Hispaniola, Montserrat, and Saint Kitts). The Jamaican population (*H. c. simulans*) is an outlier and forms a separate cluster. In a neighbor-joining (NJ) tree based on autosomal SNPs, the Jamaican population branches first from the base, suggesting it could have resulted from an earlier colonization of the Caribbean (Supplementary Figure 1). The second most basal branch includes all the mainland populations from Texas South to Peru (*H. c. bassleri* and *H. c. vasquezae*). Relationships among the remaining crown populations suggest a stepping stone model of colonization of the Caribbean islands from mainland Central America and Mexico to Cuba, thereafter following two routes: northwards to New Providence and Florida, and southwards towards the Lesser Antilles (Supplementary Figure 1).

To further explore the possibility of multiple waves of colonization of the Caribbean, we estimated admixture graphs with varying number of gene flow events. In line with the hypothesis of an earlier colonization of the Caribbean, an admixture graph with a single admixture event was best supported (Figure 1c). In this graph, the Jamaican population results from admixture between a basal lineage (44% ancestry; bootstrap confidence interval: 38-52%) and more recent lineages present in neighboring islands (56% ancestry, with confidence interval: 48-62%). The Z-chromosome retains the most variation from that earlier colonization (75.1%), compared to autosomes (25.1-40.6%; based on Twisst analysis, Supplementary Figure 2). The Jamaican population retains only the ancestral haplotypes at the mtDNA, suggesting a split at the base some 1.2 million years ago, while the extant lineages started diversifying *ca.* 670 kya. Diversification within the Caribbean began *ca.* 320 kya (based a molecular clock calibration of a Bayesian phylogenetic tree; Supplementary Figure 3).

Levels of population differentiation and diversity are also consistent with current or recent gene flow following a recent expansion and colonization of the Caribbean islands from the continent. Population differentiation, as measured by F_ST_, was generally low (mean *F_ST_* = 0.13; Supplementary Figure 4a) and absolute population pairwise divergence (*d_XY_* = 0.7% - 1.8%) is comparable to within-population nucleotide diversity (π = 0.6% - 1.5%) (Supplementary Figure 4b). Furthermore, levels of within-population nucleotide diversity (π) decrease towards the edges of the distribution and are particularly low in the island populations closest to the Lesser Antilles (Puerto Rico, Hispaniola, Montserrat, and Saint Kitts; Supplementary Figure 4b).

We crossed a female *H. c. vasquezae* from Texas and a male *H. c. tuckeri* from Florida that resulted in 32 viable adult offspring (16 females and 16 males). Both male and female hybrids were fertile: we made three F1xF1 crosses, from which 131 viable F2 individuals (67 females, 64 males) were successfully reared until eclosion. The lack of hybrid sterility is in accord with the low levels of genetic divergence across the range of the species. Hybrid female sterility is often found in other inter-species and even some intra-species crosses in *Heliconius* (Jiggins et al. 2001; Naisbit et al. 2002; Rosser et al. 2022).

We next tested the hypothesis of a recent range expansion of *H. charithonia* in two ways. First, we estimated changes in effective population size (*N_e_*) through time based both on autosomal (PSMC) and mitogenome (BSP) data. Both analyses suggest a population expansion in the recent past but at different times – *ca.* 60 kya (BSP; Supplementary Figures 5) and 200-300 kya (PSMC; Supplementary Figures 6). However, this is likely due to differences in the calibration of the molecular clock: the nuclear genome clock was estimated based on the spontaneous mutation rate between *H. melpomene* parent and offspring ^13^, while mtDNA clock was estimated based on divergence in several arthropod taxa with independently dated divergence times ^14^. Furthermore, the PSMC analysis shows that, after the initial population size increase, most populations experienced a bottleneck between *ca.* 60-100 kya, two *H. c. vasquezae* populations (Panama and Costa Rica) being the exception and showing a continued increase in *N_e_* until the recent past. Despite the initial bottleneck at around the same time as the other populations, *H. c. simulans* (Jamaica) starts expanding again *ca.* 50 kya, which could be a result of mixing between haplotypes from the more ancient and more recent colonization waves. We also used the directionality index (Ψ) ^15^ to infer the geographic origin and direction of the second wave of expansion. This test shows significant support for a range expansion (*P* << 0.001) with an origin in the range of *H. c. bassleri* in South America (Supplementary Figure 7). This is in line with phylogenetic analysis that shows *H. c. bassleri* as the most basal group of the later colonization wave (Supplementary Fig. 1). Together, all these lines of evidence suggest two colonization waves into the Caribbean from the mainland, with the Jamaican population retaining mtDNA and partial nuclear variation from the first colonization wave.

### Deeply divergent haplotypes correspond to polymorphic inversions

The lack of strong overall population structure in *H. charithonia* allows one to detect outlier genomic regions that may be under selection. We used a local PCA approach along the genome to identify genomic regions with distinct population structure. This approach has the advantage of not requiring any *a priori* definition of which population or groups of populations might be differentiated. Using this method, we found two large outlier genomic regions: one on chromosome 2 (*ca.* 1.85 Mb) and the other on chromosome 19 (*ca.* 450 kb; Figure 2a,b; Supplementary Figure 8). On chromosome 2, the PCA of the outlier region separates individuals’ genotypes into three distinct clusters (Figure 2c), the intermediate cluster having highest heterozygosity (Figure 2e). Both the intermediate and the extreme cluster with the least common genotype include only *H. c. vasquezae* individuals (from Texas and Panama); the other extreme cluster includes all other individuals. These findings are consistent with the existence of two groups of individuals homozygous for distinct non-recombining haplotypes, the intermediate cluster representing heterozygotes. The chromosome 19 outlier region yields only two very distinct clusters (Figure 2d), both groups having similar heterozygosity (Figure 2f). These likely represent homozygotes for alternative haplotypes with heterozygotes absent from our dataset. The least common genotype includes all *H. c. vasquezae* individuals from Texas, while all other individuals across the range fall into the second group. In both cases, linkage disequilibrium is high across these outlier regions when analyzing all individuals together, but not when analyzing only individuals from the most abundant cluster (Figure 2g,h). Genetic divergence (*d_XY_*) between individuals of the two extreme clusters (i.e. homozygotes) is approximately 6.32 and 6.38% within these regions, far exceeding divergence within clusters (2.14-2.74% and 1.52-2.53% for chromosome 2 and 19, respectively; Supplementary Figure 9). These patterns are consistent with large, divergent structural rearrangement polymorphisms that suppress recombination between haplotypes, allowing accumulation of divergence between haplotypes, while maintaining linkage disequilibrium along the haplotype.

**Figure 2.**
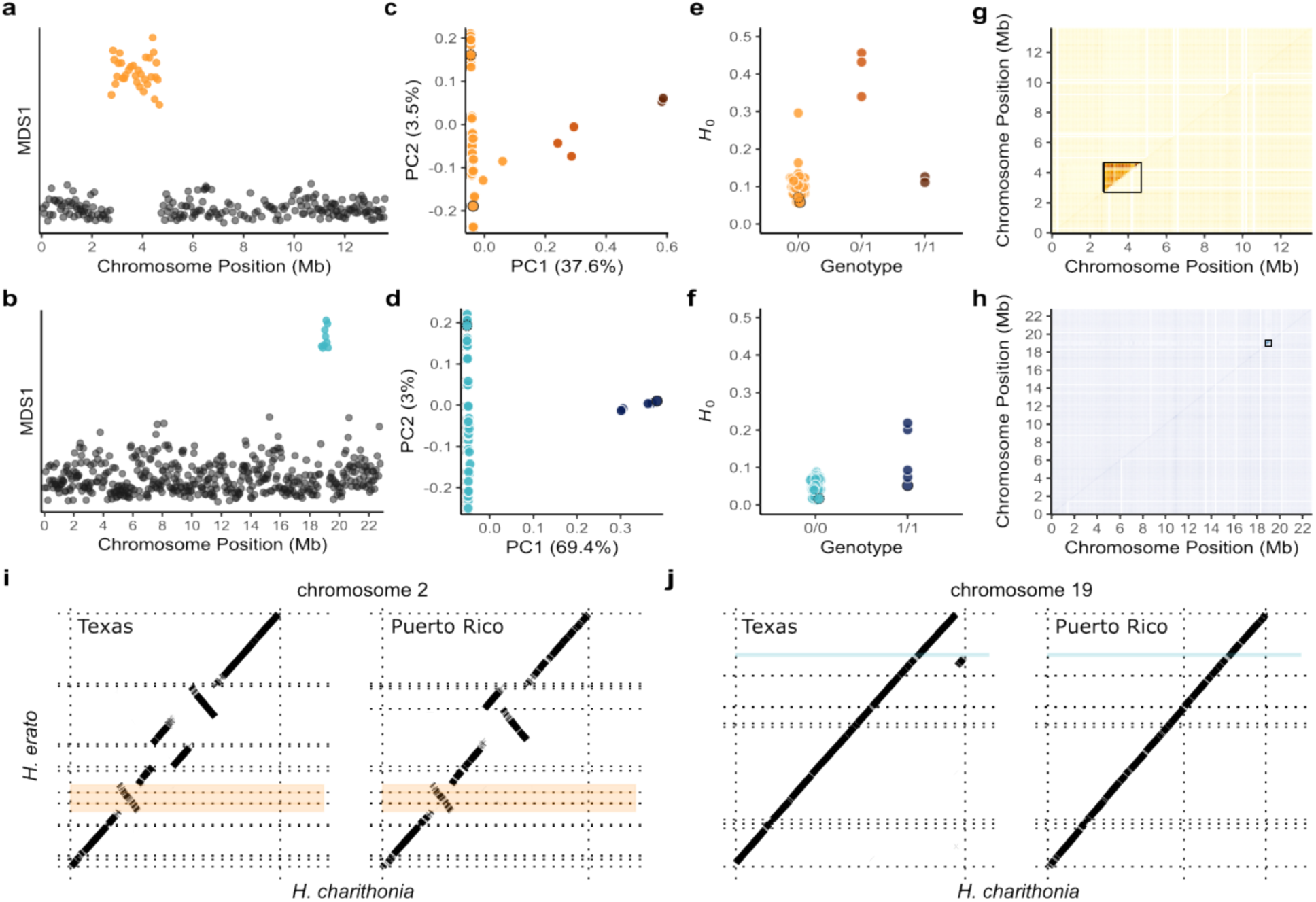
– Two large haplotype blocks appear to be inversions. (a,b) Local PCA along chromosomes 2(a) and 19 (b). Each dot represents 1000 SNP windows and windows with outlier MDS values are highlighted in orange and blue. **(c,d)** PCA of the outlier regions on chromosomes 2 (c) and 19 (d). Three and two distinct clusters are defined by principal component 1. **(e,f)** Heterozygosity at outlier regions on chromosome 2 (e) and 19 (f). Each dot represents one individual. **(g,h)** Linkage disequilibrium (LD) for chromosomes 2 (h) and 19 (g). Linkage disequilibrium was calculated including all *H. charithonia* individuals (upper triangle) or only individuals homozygous for the most common homozygous genotype (lower triangle). **(i,j)** Alignment of the Texas *H. c. vasquezae* and Puerto Rico *H. c. charithonia* assemblies to the *H. erato demophoon* reference. Only chromosomes 2 (i) and chromosome 19 (j) are shown. The putative inversions are highlighted. Dotted lines represent scaffold boundaries. These individuals are highlighted in the PCA (c,d) and heterozygosity (e,f) plots – Texas: solid black stroke; Puerto Rico: dashed black stroke.

To determine whether these divergent haplotypes are associated with structural variants we compared two genome assemblies of *H. charithonia* (from Texas and Puerto Rico) to other *Heliconius* chromosome (or near-chromosome)-level genome assemblies. Both individuals share the same genotype at the outlier region on chromosome 2, being homozygous for the most frequent haplotype in *H. charithonia* (Figure 2c,d). Genome alignments show that both individuals are homozygous for an inversion at chromosome 2 which overlaps the haploblock region, with breakpoints at positions at Herato0206:697,528 and Herato0209:424,814 (Figure 2i; Supplementary Figure 10a). Hence, the most frequent haplotype represents the inverted state of the region (absent only in some *H. c. vasquezae* individuals from Texas and Panama). At the outlier region on chromosome 19, the assemblies are homozygous for different genotypes (Figure 2d) but we found no evidence of an inversion based on genome alignments to other *Heliconius* genomes (Figure 2j, Supplementary Figure 10b). However, this is likely due to a difficulty in aligning this region due to repetitive content and/or mis-assemblies in and around putative inversion breakpoints (Supplementary Figure 11b; Figure 2j). In addition, a ∼180 kb region (Herato1910:1,521,759-1,701,968) within the putative inversion (Herato1910:1,520,293-1,973,589) aligns poorly to the *H. erato* reference genome (Figure 2j), and two highly divergent ∼ 500 kb haplotypes were assembled as separate scaffolds (Figure 2j) in the *H. charithonia* genome from Texas, one directly adjacent (upstream) of the putative inversion region and the other placed at the end of the *H. charithonia* chromosome (Figure 2j; Supplementary Figure 10b). Thus, we cannot directly show that the haploblock on chromosome 19 is an inversion, although for simplicity, we will refer to this region as an inversion and to the least frequent haplotype as the inverted haplotype.

### Evolutionary history of the inversions

To investigate the origin of the divergent haplotypes at the two inversions, we compare with closely related species from the *erato*, *clysonimus* and *sara*/*sapho* clades (Supplementary Table 1). Deep divergence between the *H. charithonia* haplotypes at the two inversions compared to genomic background levels of divergence (Supplementary Figure 9), could be explained either by ancient origin of the inversion haplotypes (i.e. retention of an ancestral polymorphism), or by introgression from other species. To test the latter, we calculated *d_XY_* along chromosomes between *H. charithonia* homozygotes and individuals representative of outgroup species. We found no drop in *d_XY_* at any of the inversion regions as would be predicted by recent introgression (Supplementary Figures 12 and 13). Maximum likelihood (ML) trees show that none of inversion haplotypes group with any other species (Figure 3). On chromosome 2, we find both standard and inverted haplotypes (the latter shared with *H. peruvianus*) are basal to the whole *erato clysonymus* + *sara/sapho* clade (Figure 3a). Also, the *H. charithonia* inverted haplotype and *H. peruvianus* group together and are deeply divergent from the rarer *H. charithonia* standard haplotype. On chromosome 19, the inverted haplotype is basal to the whole *erato* + *clysonymus* + *sara/sapho* clade (Figure 3c), while the standard haplotype retains its basal position in the *sara*/*sapho* clade as seen in the average autosomal tree (Figure 3c). The same result was also obtained when using a coalescent-aware method to estimate species trees in blocks along chromosomes (Supplementary Figure 14).

**Figure 3.**
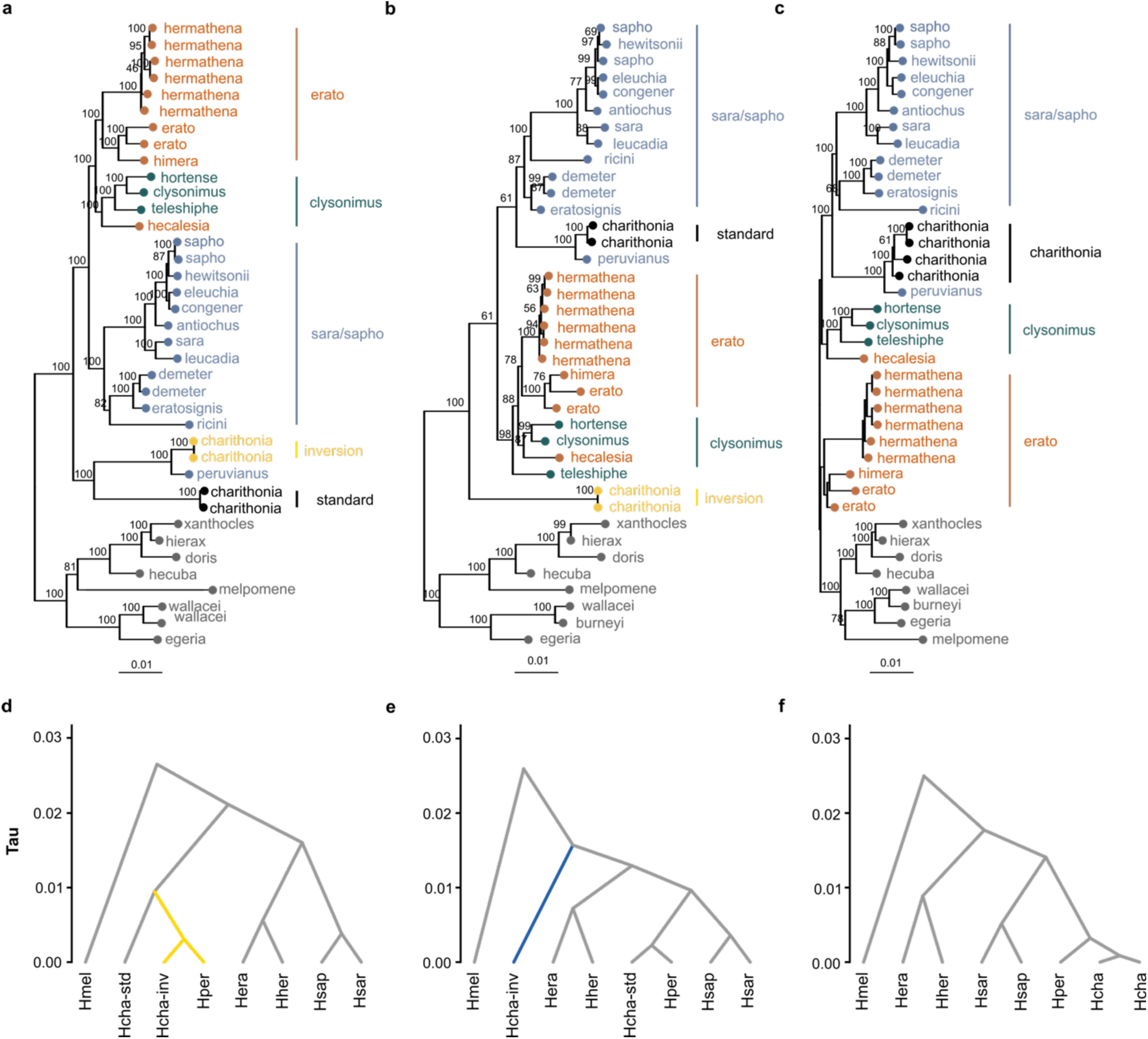
– Phylogenetics of polymorphic inversions and collinear regions of the genome. (a-c) Maximum-likelihood phylogenies of inversion regions on chromosome 2 (a) and 19 (b), and whole genome collinear regions (c). Trees were rooted using midpoint rooting. Bootstrap values indicated next to nodes. Outgroup individuals are colored according to the main clades: erato (red), sara-sapho (blue), clysonymus (green), melpomene/doris/burneyi (grey). *H. charithonia* individuals are colored according to their inversion haplotypes: inverted (gold) and standard (black). **(d-e)** Multispecies coalescent (MSC) dated phylogenies of inversion regions on chromosome 2 (d) and 19 (e) and collinear regions (f).

We estimated the ages of the two inversion polymorphisms and compared these to divergence times estimated under the majority tree from collinear regions (Figure 3d, Supplementary Table 2). The inversion on chromosome 19 originated 1.35 Mya while the inversion on chromosome 2 had a more recent origin at 816 kya. Both are more recent than the root age of *erato* + *clysonymus* + *sara/sapho* clade of 1.53 Mya estimated from collinear parts of the genome. Given their ages, and since we find neither inversion is shared with other species, the inversions appear to have been maintained as long-term polymorphisms as a result of balancing selection since their origins deep in the *H. charithonia* lineage. This could have been due either to heterozygous advantage or to local adaptation.

### Little evidence for deleterious effects of inversions

To explore possible deleterious effects of the inversions, we first investigated the inversion breakpoints. Inversions can disrupt genes if breakpoints fall within a gene or its regulatory elements ^16^. Both inversion breakpoints on chromosome 2 fall within a gene (evm.TU.Herato0206.28 and Herato0209.24, for the left and right inversion breakpoints, respectively). Their orthologs in *Drosophila melanogaster* (CG31229 and *Arc42*, respectively) are both associated with mitochondrial function. On chromosome 19, we have obtained only approximate coordinates of the inversion breakpoints based on the local PCA. The first breakpoint is within the gene Herato1910.106, an ortholog of *Catalase* in *Drosophila melanogaster*. The second inversion breakpoint occurs upstream of the evm.Herato1910.119 orthologous or paralogous to *Trehalose transporter 1-1* (*Tret1*) and *Trehalose transporter 1-2* (*Tret1l*) genes in *Drosophila melanogaster*.

We next investigated possible mutational load carried by the inversion polymorphisms, which can accumulate as a result of suppressed recombination in heterozygotes. We estimated the rate of nonsynonymous to synonymous polymorphism (pN/pS), the rate of nonsynonymous to synonymous substitution (dN/dS) and the directionality of selection (DoS), for both inverted and standard haplotypes independently. Inverted haplotypes present levels of nonsynonymous polymorphism and nonsynonymous substitution in line with the whole genome (Figure 4a,b). While levels of nonsynonymous substitution are similar to those of standard haplotypes (Figure 4b), nonsynonymous polymorphisms were significantly rarer, particularly on chromosome 19 (Figure 4a). The inverted haplotypes show overall negative (i.e. purifying) selection (DoS_chr2_ = -0.07, DoS_chr19_ = -0.01), although significantly less negative than the un- inverted haplotypes (Figure 4a). This suggests that some nonsynonymous mutations in the inverted haplotypes could be positively selected.

**Figure 4.**
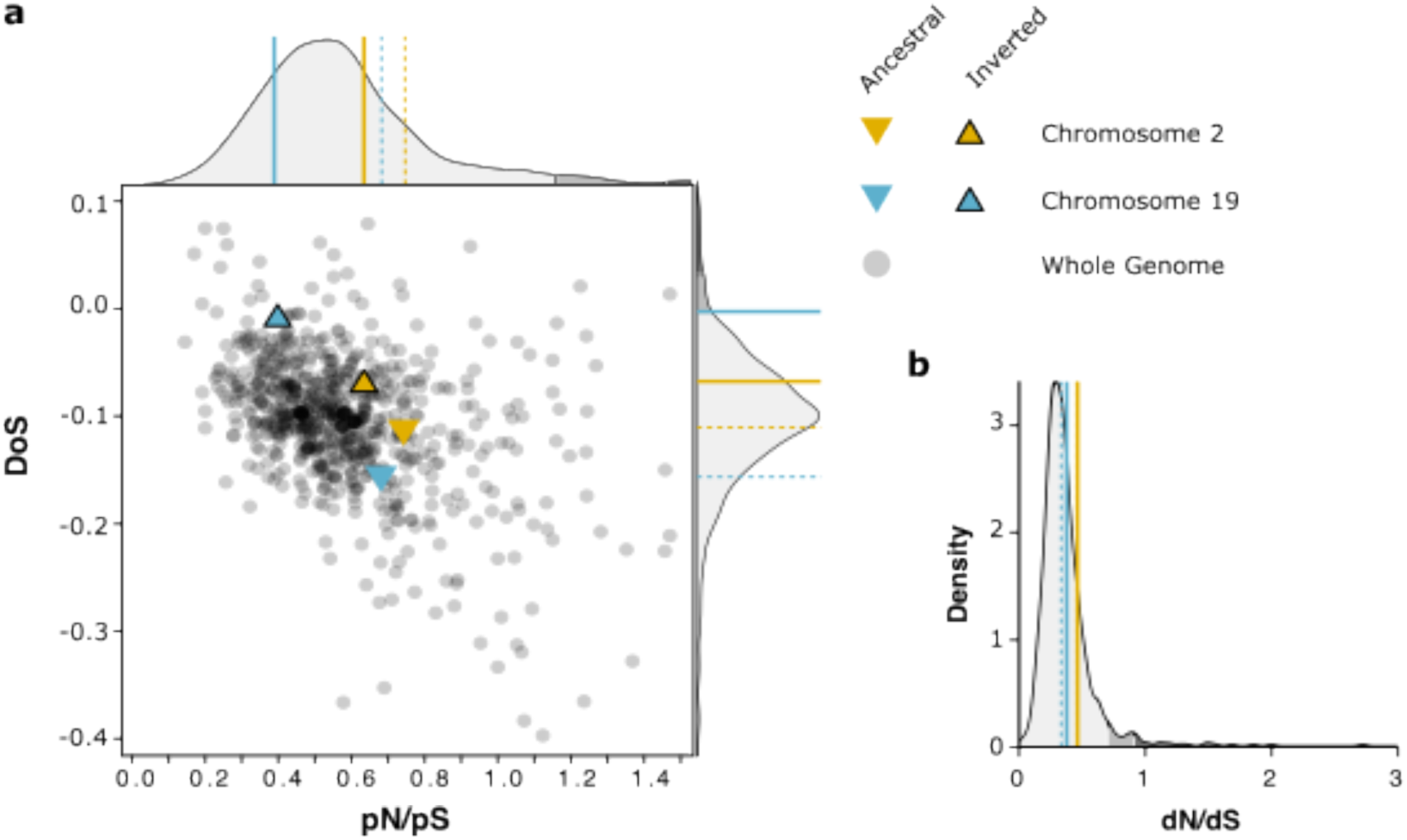
– Limited evidence for accumulation of deleterious mutations in inversions. **(a)** Direction of selection (DoS) and ratio of nonsynonymous to synonymous polymorphisms (pN/pS), computed in 500 kb windows genome-wide and in the inversion regions. **(b)** Ratios of nonsynonymous to synonymous substitutions (dN/dS). The ancestral and derived state of the inversions are given by the dashed and full colored lines. Shades of grey are used to display 0.95 and 0.975 quantiles of the genome-wide values.

If inverted haplotypes accumulate recessive mutational load, inversion homozygotes should be rare due to heterozygous advantage. Instead, heterokaryotypes were rare for the chromosome 2 inversion (n=4) and completely absent for chromosome 19 (Supplementary Table 1). Focusing only on *H. c. vasquezae* from Central America and Texas, in which the inversions are polymorphic, homokaryotypes are thus common. Chromosome 19 inversion genotype frequencies deviate from Hardy-Weinberg equilibrium in our *H. c. vasquezae* samples because inversion heterozygotes are absent. Neither inversion appears to have accumulated strongly deleterious variants, and the dearth of heterozygotes is instead consistent with homozygotes being favored by local selection, although we lacked large enough samples from any single population to test this properly.

### Inversions maintained by spatially heterogenous selection

Inversion polymorphisms may persist over the long term in a heterogeneous environment if local selection favors distinct haplotypes in different regions. Both inversions are polymorphic only in *H. c. vasquezae*. On chromosome 2, the least frequent (i.e. uninverted) haplotype is found from Panama to Texas (Figure 5a), while the rare (inverted) haplotype on chromosome 19 was only found (fixed) in our Texas sample (Figure 5b). The geographically restricted distributions of these rare inversion haplotypes are consistent with local adaptation, particularly the chromosome 19 inversion.

**Figure 5.**
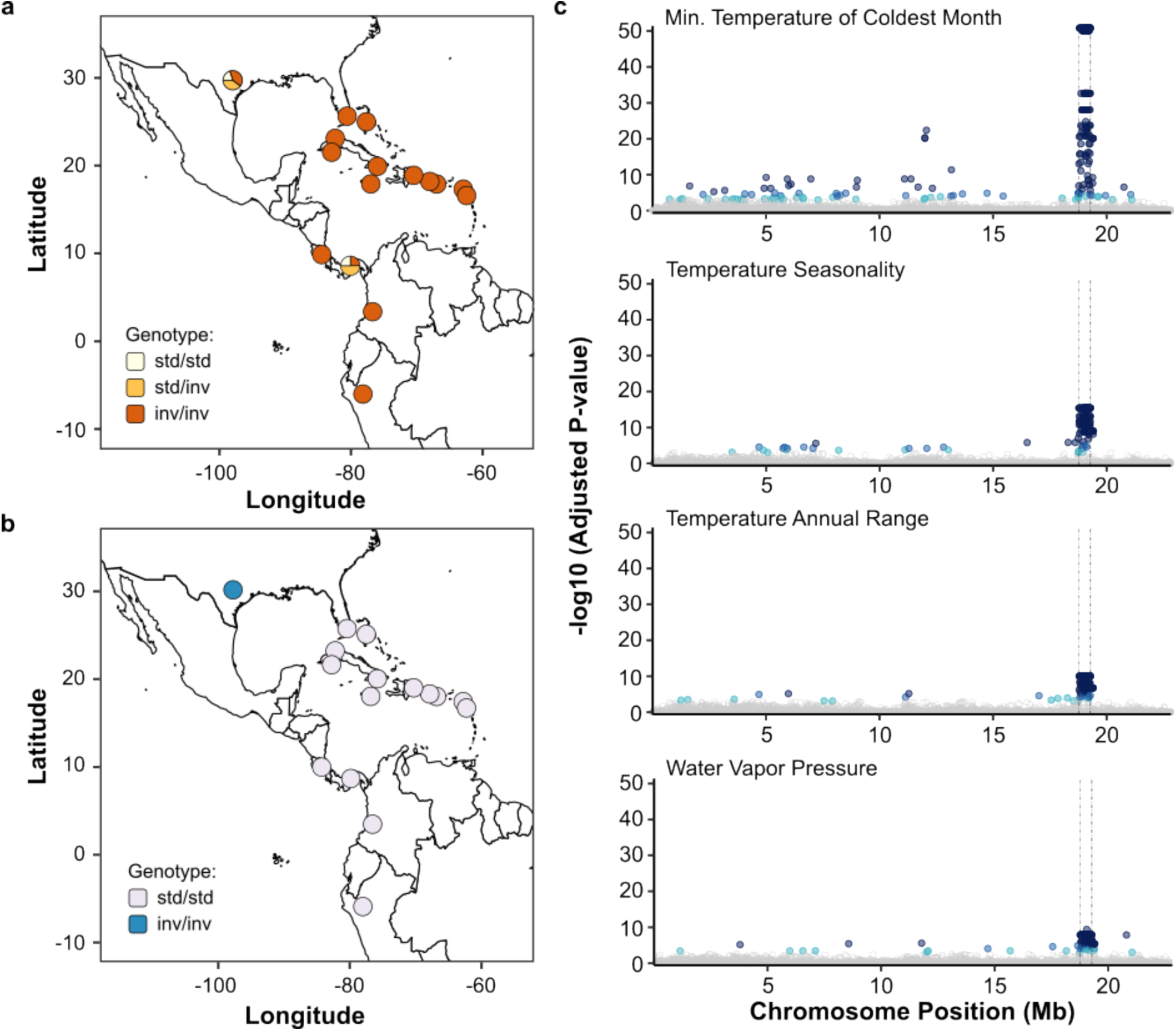
– Association of inversion haplotypes with environmental factors. (a,b) Geographic distribution of chromosome 2 (**a**) and chromosome 19 (**b**) inversion genotypes. **(c)** Latent factor mixed models Manhattan plots showing association with climatic variables inside chromosome 19 inversion. Each point indicates P-values at each SNP. Points are colored according to estimated false discovery rates (dark blue: <0.00001, medium dark blue: <0.0001, light blue: <0.001; grey: > 0.001). The dashed-dotted vertical lines represent the inferred boundaries of the inversion.

SNPs within the chromosome 19 inversion are significantly associated with four environmental variables: minimum temperature of the coldest month, temperature seasonality, temperature annual range and water vapor pressure (Figure 5c; Supplementary Figure 15). SNPs within the chromosome 2 inversion, in contrast, show no obvious association with climatic variables (Supplementary Figure 15). Populations in Texas (also representative of Northern Mexico, from which we have no samples) face significantly lower temperatures in the coldest month (3.9-7.8°C) and greater temperature oscillations throughout the year (temperature seasonality = 591-709; temperature annual range = 29-31°C), compared to *H. charithonia* from other sampled locations (temperatures in the coldest months = 7.7-22.4°C; temperature seasonality = 24-509; temperature annual range = 8-25°C; Supplementary Figure 16, Supplementary Table 3). Water vapor pressure, a proxy for humidity, is also significantly lower in Texas (0.8-1.2 kPa) than in other geographic locations (1.2-2.8 kPa). Northern Mexico similarly has long dry periods. These observations remain true even when climatic variables are sampled from locations further south in northern Mexico near the Tropic of Cancer, where *H. charithonia* overwinters before migrating north following spring and summer temperature rises (Supplementary Figure 16, Supplementary Table 3). Subtropical Texas and Northern Mexico populations thus face particularly cold and dry conditions. The rare inverted haplotype on chromosome 19 present in Texas seems likely to confer a selective advantage for these conditions.

## Discussion

Here, we present the first genome-wide analysis of *H. charithonia* population structure, with a focus on its biogeographic history and distribution of genetic diversity across its range. Using re-sequence data sampled across the species range, we infer two colonization waves of the Caribbean from the Continent into the Greater Antilles. The most recent colonization wave resulted in homogenization of genetic diversity genome-wide across the whole species range, in line with the low morphological diversity of this species. Using unsupervised methods for the detection of genomics regions with outlier genetic structure, we found two ancient large inversions segregating in *H. charithonia*, one of which likely plays a role in local adaptation to climatic conditions at the subtropical edge of its range.

### Genetic homogeneity and colonization of Caribbean islands

Unlike other *Heliconius* species with wide distributions, population structure is weak in *H. charithonia*. For example, considerable population structure exists across wide biogeographic scales in *H. erato* and *H. melpomene* ^17,18^. Deep genetic divergence found between major regions in those species is often related to major geographic barriers, for example between populations across the Andes. Within biogeographic regions there is little differentiation across most of the genome even between geographic races with different wing color patterns ^19,20^. In *H. charithonia*, these major barriers do not seem to break up genetic homogeneity, nor do we find major ecological differences between mainland and island habitats. *H. charithonia* exhibits very little genetic population structure locally ^21^, and regularly migrates hundreds of kilometers north in a single year from the Rio Grande valley on the Mexico-Texas border to Central Texas and beyond ^22^ (Fig. 1a). This long-range dispersal ability likely explains why the species is so genetically homogenous, at least on the mainland. Relatively recent, rapid expansion into the Caribbean from the continent could also help to explain the low levels of genetic divergence. We found two detectable colonization waves of the Caribbean by *H. charithonia* from our genomic dataset. The most recent colonization of the Caribbean (*∼* 320-180 kya) likely occurred from Central America into Cuba, through the Yucatan Peninsula. From there, *H. charithonia* expansion split into two routes: north into Florida and the Bahamas; and south into the Greater Antilles (Fig. 1c).

An earlier colonization (*∼*1.15 Mya) can also be inferred from genetic variation found exclusively in Jamaica, which is consistent with other studies that show this island is more isolated than other Antillean islands retaining variation from earlier Caribbean colorizations ^23^. Evidence of an ancestral colonization comes mainly from mitochondrial genomes (all four Jamaican individuals show similar deeply divergent haplotypes) and from genetic variation at the Z-chromosome (75.1% ancestral variation). In contrast, the autosomes retained much less ancestral variation (31.4%). The contrast between autosomal and mitochondrial loci was suggested to result from male-biased migration into the Caribbean and/or different effects of founder events and genetic drift on the different genomic compartments ^12^. Our data from the Z-chromosome allows us to test between the two hypotheses. In ZW systems such as *H. charithonia*, male-biased dispersal would favor an excess, not a deficit of homogenization of variation on the Z-chromosome (males are ZZ). Instead, the ancestral variation in different compartments of the genome is more consistent with differential impact of founder events and genetic drift.

### Evidence for chromosomal inversions

The availability of population-level whole-genome datasets allow the discovery of structural variants in a comprehensive and cost-effective way ^24^. For example, an increasing number of studies have investigated genetic heterogeneity along the genome using unsupervised methods such as PCA in sliding-windows ^25^. Such methods have revealed genomic blocks of tightly linked SNPs, which are highly differentiated, providing clues to structural variants such as chromosomal inversions that reduce recombination ^26–31^. Complementary analysis, including PCA within these showing three distinct genetic clusters and elevated heterozygosity of the putative heterokaryotypes, further supports the presence of an inversion polymorphism. This unsupervised and systematic approach to detection of structural variants has two advantages. First, since potentially diverging groups do not need to be defined *a priori*, it does not limit the number of axes of differentiation that can be explored. Second, because inversions are first characterized before testing adaptation, it can provide an unbiased test for the role of inversions in adaptation. However, uncovering inversions from indirect evidence has its own drawbacks. Inversions are best confirmed using direct methods such as long read sequencing and/or HiC sequencing data ^32,33^. Direct methods are also better for characterizing inversion breakpoints, which can only be approximated using indirect approaches. Also, because indirect methods rely on signals of linkage and divergence, they can most easily detect large, polymorphic and highly divergent inversions ^34^.

Here, we employed such an unsupervised approach to discover two highly differentiated genomic regions which are very likely inversions. The first inversion is *ca.* 1.4 Mb long and falls on chromosome 2. Curiously, several other chromosome 2 inversions have been described previously in other *erato-clysonimus-sara/sapho* species ^31,35–37;^ however, the inversion we found in *H. charithonia* is not homologous to any of these. The second inversion, on chromosome 19, is ca. 450 kb long and is also not previously described from *Heliconius*. We were able to directly confirm the inversion on chromosome 2 and determine its breakpoints, based on genome alignments to outgroup species, but we were unable to do so for the putative inversion on chromosome 19. The two *H. charithonia* assemblies (from Puerto Rico and Texas) available have distinct genotypes at this chromosome 19 region (both homozygous, but with different inversion haplotypes), although genome-genome alignments show them both to be collinear to outgroup species. While it is possible that this region does not correspond to an inversion, we believe that we were unable to confirm it due to a mis-assembly in one of the reference genomes, probably the Texas assembly (i.e. the inversion was not assembled in the correct orientation). Two lines of evidence support this hypothesis. First, coverage is higher near the boundaries of the chromosome 19 region compared to genome average, suggesting the breakpoints might be rich in repetitive sequences (Supp. Figure 11). Genome assembly across repetitive regions can be challenging, particularly when generated from short-read data. This is the case of the Puerto Rico assembly, which was generated based on linked short-read data (10X Chromium technology). The Texas assembly was, however, generated using PacBio and Hi-C data. Provided the reads were large enough to span the repetitive regions around inversion breakpoints, this assembly should have the putative inversion region in the correct orientation. However, when we align the *H. charithonia* genome from Texas to outgroup species, we find that one *ca.* 500 kb region directly adjacent and upstream of the putative inversion is apparently duplicated as it is also present at the end of chromosome 19. When mapping this individual’s PacBio reads to its own genome assembly we find coverage at these regions drops to half of genome-wide coverage (Supplementary Figure 18). This suggests that these represent two distinct haplotypes of the same genomic region. However, due to high divergence between the two, they were assembled separately, with one being placed at the end of chromosome 19 (Supplementary Figure 18). This underscores the complex genomic structure around the putative inversion region, and we cannot discard the hypothesis that this region was misassembled also.

### Ancient polymorphic inversions

Chromosomal inversion polymorphisms are often ancient and can pre-date the origin of the species in which they are discovered ^38^, which is sometimes explained by introgression from closely related species ^39–42^, including in *Heliconius* ^31,35,36,43^. We find that inversions segregating in *H. charithonia* are old, both pre- dating the split between *H. charithonia* and its sister species *H. peruvianus* but found no evidence of introgression. While we cannot discard past introgression from an extinct or unsampled species, it seems more likely these inversions are ancient ancestral polymorphisms. In fact, the phylogenies of both inversion regions show that the inversion haplotypes are basal to the entire *erato-clysonimus-sara/sapho* clade (Figure 3), suggesting they pre-date the origin of the group. In the case of the inversion on chromosome 2, this is unlikely since the origin of the inversion haplotype (816 kya) dates more recently than the root age of the erato-clysonimus-sara/sapho clade (1.54 Mya) inferred from collinear regions of the genome. Instead, the date of the inversion and its topology can be reconciled if we assume introgression from the *erato* or *clysonymus* clade into the *sara/sapho* clade, after the split of the latter group from *H. peruvianus* and *H. charithonia*. Regarding the putative inversion on chromosome 19, the posterior mean estimate for the age of the inversion (1.35 Mya) is slightly younger than the root age of the erato-clysonimus-sara/sapho clade (1.53 Mya), though their confidence intervals overlap (Supplementary Table 2). However, node ages estimated from collinear parts of the genome (including root age) are likely underestimated since our analysis does not consider gene flow, which is common in this clade ^44^. As a result, it remains plausible that the chromosome 19 inversion predates the origin of the entire group.

The long-term persistence of inversion polymorphisms raises questions about how inversions become established and maintained within species. Multiple factors – including local adaptation, mutation load, breakpoint effects – have been suggested to explain these processes and their relative role may change through the life of an inversion ^5,9,45^. For instance, in *Heliconius numata*, wing-color polymorphisms is associated with an inversion at a mimicry locus (supergene P) on chromosome 15 ^46,47^. In this system, the Müllerian mimicry adaptations associated with recombination suppression are thought to explain the initial spread of the inversion polymorphisms, but this may be coupled with assortative mating among phenotypes and accumulation of deleterious mutations leading to heterozygous advantage ^48^.

In *H. charithonia*, there is little evidence for accumulation of mutational load in the form of non-synonymous mutations in either inversion. In the absence of recombination, inversions that are frequently heterozygous are expected to accumulate deleterious mutations, and each chromosomal arrangement may be fixed for different mutations, leading to greater fitness of heterozygotes. Recombination is suppressed only in heterokaryotypes but can proceed in homokaryotypes, so that purifying selection can remove deleterious mutations when both chromosomal morphs have reasonably large effective population sizes ^49^. In *H. charithonia*, chromosome 2 heterokaryotypes were rare (n=4) species-wide (n=55), and completely absent for the chromosome 19 inversion (Figure 2f; Supplementary Table 1). Even if we consider only *H. c. vasquezae*, the subspecies with the chromosome 2 polymorphism, inversion genotypes show no evidence of heterozygous excess, suggesting that neither inversion haplotype has fixed, highly deleterious mutations. A perhaps more plausible explanation for the establishment and maintenance of inversion polymorphisms in *H. charithonia* is local adaptation. Inversions can be then maintained as a stable polymorphism in a heterogenous geographical landscape by migration-selection balance ^5,50^. If a new inversion happens to capture a set of locally adapted alleles at two or more loci, the inverted haplotype will be advantageous, since suppressed recombination within the inversion preserves the favorable combination of alleles ^5^. The spatial structure of the chromosome 19 polymorphism and its association with environmental conditions (Figure 5a,c), suggest that different inversion haplotypes confer local benefits in response to climatic conditions. The weak geographic structure at the whole genome level indicates that populations are currently connected by high levels of gene flow, which, in the inversion region, is counteracted by divergent selection. Notably, the inversion haplotypes present in Texas resisted a recent population expansion that homogenized most genetic variation across the entire species range. While it is not clear whether this inversion was initially advantageous and spread because of recombination suppression, it is likely currently maintained by a balance between migration and selection.

### Chromosomal inversions involved in adaptation to heterogenous environments

Clinal variation of inversions across geographical or environmental gradients offers compelling evidence of natural selection driven by abiotic factors ^38,51^. Numerous recent examples of species in which chromosomal inversions segregate between distinct ecotypes or in parallel with environmental gradients, include monkeyflowers ^52^, sunflowers ^26,27^, deer mice ^28^, seaweed flies ^29^, annual ragweeds ^30^, marine snails ^53,54^, cod ^55^, sticklebacks ^56^, and *Heliconius* butterflies ^31,37^. Likewise, in *H. charithonia* the two inversion polymorphisms exhibit geographic structuring, suggesting a potential role in local adaptation.

Different lines of evidence indicate that the inversion on chromosome 19 confers a local adaptive advantage, particularly in response to cold temperatures and desiccation stress. First, genotype-environment analyses show an association between a cluster of SNPs within the chromosome 19 inversion and climatic variables related to temperature variability throughout the year, cold and desiccation (Figure 5c; Supplementary Figure 15). A previous study also found that temperature was a key factor determining *H. charithonia* length of residency time in Texas during the warmer months of the year ^22^. Second, the inversion polymorphism appears to be spatially segregated between populations in dry and cold habitats (restricted to in Texas) and populations in warmer and more humid habitats (elsewhere in the rest of the *H. charithonia* distribution; Figure 5b, Supplementary Figure 17). Thirdly, the chromosome 19 inversion contains 16 genes, including two located close to the inversion boundaries – *Catalase* (*Cat*) and *Trehalose transporter 1*-*like* (*Tret1l*) –, that have been implicated in adaptation to cold and desiccation across different taxa ^57–61^. In insects, species resistant to cold also tend to be tolerant of desiccation ^62,63^, and many mechanisms of tolerance to these two environmental stresses overlap ^62^. Among others, these include upregulation of antioxidant defenses and metabolism and transport of trehalose between the cells and the hemolymph. Periods of environmental stress, such as cold and desiccation, result in an increase of reactive oxygen species (ROS) which can cause cell damage ^63,64^. In response, increased activity or expression of antioxidant enzymes helps mitigate oxidative stress, including Catalase which breaks down harmful hydrogen peroxide into water and oxygen ^57,58,65^. Another key response to extreme conditions, such as cold, heat, desiccation, is biosynthesis and transport of sugars, especially trehalose. Trehalose is the main hemolymph sugar in most insects and acts as a cryoprotectant at low temperatures. Accumulation of trehalose improves tolerance to cold, desiccation, and hypoxia and facilitates cryoprotective dehydration in insects by replacing water and preserving the structures of proteins and membranes during stress ^66–69^. Given its geographic structure, association with environmental variables and known functions of genes within, it is highly likely that the chromosome 19 inversion polymorphism is involved in adaptation to drier and colder subtropical conditions in the northern part of the *H. c. vasquezae* range, enabling its success as the most northerly distributed member of its genus.

The inversion polymorphism on chromosome 2 showed no association with any of the climatic variables we examined. However, a GO enrichment analysis revealed an overrepresentation of genes associated with ‘wing disc’ and ‘gustatory receptor neuron’ phenotype categories within this inversion. The latter phenotype suggests the chromosome 2 inversion may play a role in host plant adaptation. Notably, *H. charithonia* is among the few *Heliconiini* adapted to and able to feed on Passiflora hosts with hooked trichomes ^70,71^. This adaptation is also variable within *H. charithonia*: larvae from mainland populations (*H. c. vasquezae*) can escape entrapment and physical damage from the trichomes by laying silk mats on the trichomes ^70,71^; in contrast, larvae from the island of Puerto Rico, where hooked trichome *Passiflora* do not occur, become entrapped by the hooked trichomes of Central American species (W.O.M., personal observations in insectaries). Coincidently, the distribution of *Passiflora* species with hooked trichomes, such as *Passiflora adenopoda* and *Passiflora lobata*, is restricted to northern South America and Central America. These *Passiflora* distributions overlap with the range of *H. c. vasquezae* (Central America), in which both inversion haplotypes are present, but also extends to that of *H. c. bassleri* (South America) which is fixed for the same uninverted haplotype found on the island populations. Different reasons could explain the lack of a perfect association between the ranges of inversions haplotypes and the Passiflora with hooked trichomes: the relative densities of Passiflora with and without trichomes (which exist in these areas) may change through Central and South America; we might have missed inversion haplotypes in *H. c. bassleri* due to the low sample size (n=4). However, a true lack of association between chromosome 2 inversion and host plants must also be considered.

Experimental assays and genetic mapping will be fundamental to establish a more direct test of the involvement of inversion haplotypes and putative adaptive traits, as demonstrated in other species ^29,72,73^. This should be particularly feasible in this system, given the low genomic divergence outside the inversion regions and the full fertility observed between subspecies.

## Methods

### Sample collection and genome re-sequencing

We performed Illumina short-read whole genome sequencing of 47 *H. charithonia* collected from across most of *H. charithonia* native range (Supplementary Table 1, Figure 1a). These included wild-caught specimens collected from different localities across the species ranges between 1990-1991, wild-caught samples collected in Peru in 2011, and fresh samples reared in captivity from Colombia, Costa Rica and Texas (Supplementary Table 1). RNA-free genomic DNA was extracted using the E.Z.N.A Tissue DNA kit (Omega Bio-tek, Inc.), including an RNase A treatment step. DNA integrity was manually inspected on agarose gels and concentrations were determined on a Nanodrop Spectrophotometer. Whole genome DNA library preparation was performed using the Illumina DNA Prep library kits aiming at an insert size of ∼350 bp. The resulting libraries were sequenced using 150 bp paired-end sequencing on Illumina NovaSeq S4 and SP instruments at the Harvard University Bauer Core (see Supplementary Table 1). To our new data we added previous whole genome re-sequence data of seven *H. charithonia* individuals, and 28 genomes of 18 closely related species (Supplementary Table 1).

### Read mapping and genotype calling

Reads were filtered for adapters using Trimmomatic v0.39 4 and mapped to the *H. erato demophoon* reference genome 5 using bwa-mem v0.7.15 ^76^ with default parameters. Median coverage across all *H. charithonia* samples was 11.2X (ranging from 4.6X to 126.0X; Supplementary Table 1). Genotyping was carried out with bcftools v1.17 *mpileup* and *call modules*, using the multiallelic-caller model (call -m), requiring minimum base and mapping qualities of 20. Genotypes were filtered using the bcftools *filter* module. Both invariant and variant sites were required to have a minimum quality score (QUAL) of 20. Furthermore, individual genotypes were filtered to have a depth of coverage (DP) ≥ 4 (except for the Z-chromosome of females for which the minimum required DP ≥ 2) and QUAL ≥ 20. Genotypes not fulfilling these requirements or within 5 bp of an indel (–SnpGap) were recoded as missing.

### Population Structure and Phylogenetic Analysis

Population structure was investigated using a principal component analysis (PCA) as implemented in PLINK v1.9 7. We considered only *H. charithonia* individuals and only biallelic sites (excluding singletons) with no missing genotypes, each at least 25 kb from the next SNP. This was done separately for the autosomes and the Z-chromosome (13,306 and 610 independent SNPs, respectively). The same datasets were further used to examine the degree of shared genetic variation between different samples using fastSTRUCTURE ^78^. We ran fastSTRUCTURE for *K* ranging from 1 to 5, and used the chooseK.py script to choose the best value of *K*.

We estimated phylogenies all *H. charithonia* genome sequences and one of the sister species, *H. peruvianus*. Autosomal and Z-chromosome phylogenies were generated separately, using both variable and invariable sites. In PLINK v1.9 ^77^, positions were selected to be at least 1 kb apart (--bp-space 1000) with no missing genotypes (--geno 0) across all 54 *H. charithonia* individuals and the *H. peruvianus* outgroup. The resulting vcf file was converted to FASTA format using a custom script (https://github.com/FernandoSeixas/Hcharithonia-inversions). We inferred neighbor-joining (NJ) trees in the phangorn R package ^79^. Pairwise distances accounting for saturation were first calculated assuming the Jukes-Cantor (JC69) model of DNA evolution. Neighbor-joining phylogenies were then estimated using the NJ function, using the ‘dist.ml’ and ‘NJ’ functions, respectively. The phylogenies were midpoint rooted.

To infer population relationships while accounting for admixture events, we estimated admixture graphs, using the ‘*qpgraph’* function of the ADMIXTOOLS 2.0.0 R package ^80^. Only autosomal SNPs (excluding singletons), with no missing genotypes (--geno 0) and at least 1 kb apart (--bp-space 1000) were considered, resulting in 255,569 SNPs. We ran the analysis grouping individuals by their assigned population (see Supplementary Table 1). We considered admixture graphs with up to four admixture events, each estimated using three independent replicate runs. In each run, the best-fit graph was estimated using the command ‘*find_graphs’* with default parameters and specifying *H. peruvianus* as outgroup. For each number of admixture events only the best run (i.e., the run with the lowest score) was considered. We then determined which model was best supported, by running 100 block-bootstrap replicates of the best graph under each model and comparing the likelihood score distributions using the ‘*qpgraph_resample_multi*’ and ‘*compare_fits*’ functions. Confidence intervals for strength of drift and admixture proportion were estimated using the ‘*qpgraph_resample_snps*’, with 100 bootstraps.

We estimated a dated phylogeny for the mitochondrial genome. Whole mitochondrial genome sequences of each individual were assembled from a subset of 5 million trimmed reads with MITObim v1.9.1 ^81^, using the –quick option and up to 40 iterations. The full mitochondrial genome of *H. sara* was used as bait (Genbank accession NC_026564). Mitochondrial genome assemblies were aligned using MAFFT ^82^, and pruned manually in Geneious version 2023.2.1 ^83^. We selected only genic regions for phylogenetic analysis, based on annotations of the *H. sara* reference. Models of DNA evolution for each gene alignment were fit using ModelTest-NG ^84,85^. Bayesian phylogenetic inference in BEAST v2.6.3 ^86^ was used to date divergence times. Three independent runs of 10 million generations were performed using the best-fit nucleotide substitution model (or the next-most simple model implemented in the software), a Bayesian Skyline Plot tree prior, and a strict molecular clock. Runs were examined in Tracer v1.7.1 ^87^ for convergence and consistency across runs. Replicate runs were concatenated using LogCombiner and post-burn-in trees were summarized using TreeAnnotator, both part of the BEAST package. Node ages were calibrated assuming a substitution rate of 1.15 x 10^-8^ substitutions/site/year for the COI region ^88^. We also constructed a median-joining network using PopART 1.7 ^89,90^.

### Summary statistics and genomic differentiation

Within-population (π) and between-population (*d*_XY_, Hudson’s *F_ST_*) population summary statistics were estimated in sliding windows of 50 kb (50 kb step) along the genome using the python script popgenWindows.py (available from github.com/simonhmartin/genomics_general). Sites with less than 80% individuals genotyped were discarded; only windows with at least 20% sites passing filters were considered. We found that the Jamaican population retains variation from an ancestral colonization wave of the Caribbean. To explore heterogeneity in ancestry across the genome we used Twisst ^91^ (https://github.com/simonhmartin/Twisst). Phylogenetic relationships were estimated among three focal subspecies (*H. c. simulans* (Jamaica), *H. c. churchii* (Dominican Republic), and *H. c.* bassleri (Colombia and Peru)), using *Heliconius peruvianus* as the outgroup. Only SNPs variable in the focal species and lacking missing data were considered. Statistical phasing and imputation were performed using Beagle v5.1 ^92^, with default settings. Neighbor-joining trees were inferred from the phased filtered dataset, in 50 kb non-overlapping windows, assuming a GTR substitution model, in PHYML ^93^. Exact weightings were computed for all phylogenies.

### Genetic crosses

Captive-bred populations of *H. charithonia* from Florida and Texas were reared in the OEB Harvard greenhouses. Adult butterflies were fed with a solution of water with sugar and pollen and provided additional pollen sources – *Lantana* spp. (Verbenaceae). We performed crosses between captive bred *H. c. vasquezae* (Texas) with *H. c. tuckeri* (Florida). F1 individuals were obtained by crossing one pure Texas virgin-female with one pure Florida male. Three F1 female-male pairs were mated to generate F2 progeny. Females were kept isolated from males prior to the crosses to ensure all were unmated. Individuals were monitored daily and stored once adult butterflies emerged.

### Historical demography and range expansion

Past demographic dynamics of *H. charithonia* were estimated using the Pairwise Sequentially Markovian Coalescent (PSMC) model ^94^. Diploid consensus sequences were obtained using samtools v1.17 *mpileup* for all autosomal contiguous scaffolds longer than 1 Mb and requiring a minimum base and mapping quality of 20 and depth of coverage ≥ 8. Because of PSMC’s limitations inferring coalescent rates in the more recent past, we also used coalescent Bayesian Skyline Plot (BSP) ^95^ in BEAST v2.6.3 14 for the mitochondrial genome, as described above. Since the *H. charithonia* mitochondrial lineage from Jamaica represents a basal and divergent lineage, the Jamaican mitogenomes were excluded from this analysis. The Bayesian Skyline demographic profile was generated in Tracer v1.7.1 15 and plotted with R. e tested for range expansion of *H. charithonia* vs. equilibrium isolation-by-distance using the method outlined in ^15^. This method relies on allele frequency clines created by successive founder events during a range expansion and can infer the strength of the founder effects associated with spatial expansion and the most likely expansion origin ^15,96^. The data was prepared using PLINK v1.9 ^97^, considering only SNPs with no missing data and, a minimum allele frequency of 0.05 and at least 10 kb apart. The *H. peruvianus* individual was included to determine the derived state of each allele. The filtered dataset was then analyzed in the R package “rangeexpansion”. This analysis was performed both with all individuals or excluding individuals from Jamaica.

### Detection of divergent haplotypes

#### Local PCA

To identify genomic regions with outlier population structure, we performed local principal component analysis (PCA) with the *lostruct* R package ^98^. We used the dataset including only *H. charithonia* with biallelic sites (excluding singletons) with a maximum of 5% missing genotypes. Local PCA in *lostruct* was performed for non-overlapping windows of 500 SNPs and independently for each chromosome using the *eigen_windows* function. The distance matrix between windows from local PCs was then computed using the *pc_dist* function (with the two top PCs) with default parameters and distances were visualized using multi-dimensional scaling (MDS) with the *cmdscale* function with two MDS axes. The two MDS axis were then visualized by plotting the MDS score against the genomic position of each window. The z-score of the MDS1 score for each window was calculated; potential haploblocks of interest corresponded to genomic regions with at least 5 consecutive windows with a z-score > 3.

#### PCA and heterozygosity

All SNPs within these haploblocks were used to calculate PCAs using PLINK 1.9 ^77^. The k-means algorithm from the *kmeans* package in R was used to define clusters from PC1. Given the low sample size of some clusters, we followed the same approach as in ^99^, and defined the starting positions for each cluster as the minimum, maximum and middle values of PC1 scores. For each region, we also measured heterozygosity (π) in vcftools ^100^.

#### Linkage disequilibrium

For each chromosome harboring a putative inversion, we estimated pairwise LD (*r*^2^) considering either 1) all *H. charithonia* individuals or 2) only individuals of the most represented cluster as defined by the PCA of the region. In this analysis we excluded two individuals with more than 20% missing data. Only bi-allelic SNPs with MAF > 0.05 and at least 1000 bp apart were considered. Genotype *r*^2^ values were calculated with vcftools geno-r2 ^100^. Finally, the mean *r*^2^ values were calculated between all 100 kb windows within a chromosome were calculated using the script emerald2windowldcounts.pl (from https://github.com/owensgl/reformat).

#### Genetic differentiation

To measure genetic differentiation between standard and inversion haplotypes we calculated absolute sequence divergence (*d*_XY_) using the python script popgenWindows.py (available from github.com/simonhmartin/genomics_general). This was calculated in sliding windows of 50 kb (50 kb step) along chromosomes harboring the inversions, between the predicted homozygote genotypes.

#### Comparison of genome assemblies

To determine whether divergent haplotypes correspond to inversions we compared the chromosome level genome assemblies of one *H. c. vazquezae* individual from Texas ^101^ and one *H. c. charithonia* from Puerto Rico ^102^ to the two *H. erato* ^75,103^, the *H. sara* ^104^ and the *H. melpomene* Hmel2.5^105^ reference genomes. Identification of structural variants was performed using SyRI 1.6.3 ^106^. SyRI expects chromosome-level assemblies with the same number of chromosomes. Hence, for the genome assemblies of *H. melpomene*, *H. erato demophoon, H. e. lativitta* and *H. charithonia* from Puerto Rico, we concatenated scaffolds in the same chromosome. Pairwise genome to genome alignments were them performed using minimap2 ^107^, with parameters ‘-ax asm20 --eqx’ and structural variants were then identified using SyRI, with parameters ‘-f -k’.

### Timing the origins of inversions

To reconstruct the evolutionary history of the two inversions, we analyzed an extended dataset including 8 closely related species (26 subspecies; Supplementary Table 1). Read mapping and genotype calling were performed as described above.

#### Summary Statistics

Absolute genetic distance (*d_XY_*) between both the standard and inverted haplotypes and the outgroups was calculated in sliding windows of 50 kb (50 kb step) along chromosomes using the python script popgenWindows.py (available from github.com/simonhmartin/genomics_general).

#### Phylogenetic analyses

Phylogenetic relationships within the inversion regions were estimated based on maximum likelihood (ML) concatenated gene trees using IQ-TREE (v2.1.0) ^108^. Two *H. charithonia* individuals, homozygous for each inversion haplotype, and individuals representative of the different *Heliconius* clades were included. Sites without missing information in all individuals were considered. Model selection was performed using ModelFinder ^109^ (-m MFP) and branch support was estimated using ultrafast bootstrap implemented in IQ-TREE ^110^, with 5,000 ultrafast bootstrap replicates (-B 5000). To retain information at heterozygous sites, we assigned IUPAC ambiguity codes to which IQ-TREE assigned equal likelihood for each underlying base identity.

Phylogenetic relationships across the genome were also estimated for *H. charithonia* harboring different inversion genotypes, as well as outgroup species, using the multispecies coalescent (MSC) approach implemented in BPP v.4.6.2 ^111^ Only a subset of species was considered, and *H. melpomene* was used as an outgroup. Loci were selected to be 300 bp long, at least 2 kb apart from the nearest loci and at least 2kb apart from exons as annotated in the reference genome. Repetitive elements as annotated in the reference genome were masked before producing sequence alignments. For each locus, individuals with more than 50% missing genotype calls were excluded from the alignment and only loci with at least two individuals per population were considered. Furthermore, sites with more than 20% of individuals with missing genotype calls were removed and loci with less than 50 bp passing filters were excluded. Loci were grouped into blocks of 100 loci, and those overlapping the inversions on chromosomes 2 and 19 were grouped in separate blocks. Species-tree estimation was then performed in BPP v.4.6.2 using the A01 analysis (species-tree inference assuming no gene flow). Inverse gamma priors (invGs) were applied both to the root age (τ0) and to effective population sizes (θ) – invG(3, 0.3) and invG(3, 0.04), respectively. Parameters were scaled assuming a mutation rate of 2.9 × 10^−9^ substitutions per site per generation and a generation time of 0.25 years ^13^. The MCMC was run for 1,000,000 iterations after 32,000 iterations of burn-in, sampling every 2 iterations.

#### Dating of inversions

To estimate divergence times between standard and inversion haplotypes we again used BPP, but assuming a fixed species tree (A00 analysis). For each inversion, the inversion trees inferred from the BPP A01 analyses were used. We also estimated divergence times between species, using the collinear parts of the genome. For this, we assumed the majority tree across all non-inversion blocks to be the true species tree. To reduce the amount of data, we subsampled 10 loci from each non-inverted blocks in chromosome 2 and 19. The analysis were run using the same priors, burn-in and number of iterations as in the BPP A01 analyses.

### Functional impact of inversions

#### Genes near inversions breakpoints

We used the *H. erato demophoon* gene annotation ^112^ to explore whether inversion breakpoints disrupted annotated genes. The coordinates of inversion breakpoints on chromosome 2 were determined based on the genome alignments, while for the chromosome 19 we relied on the local PCA results. For each inversion we recorded genes spanning the breakpoints or the closest annotated gene to the left and right of the inversion breakpoint.

#### Mutational load

To test whether inversions are enriched for deleterious mutations, we calculated the ratio of synonymous to nonsynonymous polymorphisms (pS/pN) within *H. charithonia*, the ratio of synonymous to nonsynonymous substitutions (dN/dS) compared with *H. e. demophoon*, and the direction of selection (DoS) ^113^. SNPs in *H. charithonia* were annotated using SNPEff v5.1d ^114^, with default parameters, and the *H. erato demophoon* reference genome annotation. To ensure each gene comprises several SNPs, only genes larger than 5 kb were considered. These metrics were calculated for each inversion region using only individuals homozygous for each of the inversion haplotypes, while whole genome distributions were obtained using all individuals and calculated on 500 kb non-overlapping windows.

#### Hardy-Weinberg equilibrium (HWE)

Deviations from HWE within *H. c. vasquezae*, for inversion genotype frequencies was using HWE.chisq in R from the ‘*genetics*’ package.

#### Environmental Associations

To identify SNPs associated with environmental variables, we used latent factor mixed models, as implemented in LEA R package ^115–117^. This analysis tests for significant associations between SNP allele frequencies and the selected environmental variables after correcting for genetic structure. SNPs were filtered to include only those with no missing data, a minimum allele frequency (MAF) of 0.05 and at least 100 bp apart from each other, resulting in 3,384,859 SNPs. The climatic variables were obtained from the WordClim database, with 5 arc-minutes (ca. 86 km^2^) resolution (Hijmans et al., 2005), and extracted for each specimens’ location using QGIS 3.36.2. Genotype-environment associations were inspected using the lfmm2 function, with the lambda default parameter of 1 x 10^-5^, and number of factors K = 2. The resulting p-values were adjusted to account for multiple testing using the Benjamini-Hochberg method.

### Permits

The export of DNA extractions from samples stored at STRI, Panama, was approved local authorities, the Ministerio de Ambiente – Dirección de Áreas Protejidas y Biodiversidade, Sección de Acceso a Recursos Genéticos y Biológicos (SARGEB), permit number PA-01-ARG-046-2022.

## Acknowledgements

We thank the following: the Harvard FAS Research Computing team for their support. Hopi Hoekstra (Harvard University) for laboratory space. Nick Grishin (UT Southwestern) for providing whole genome sequencing data from Texas and Florida individuals. Adriana Briscoe (UC Irvine) and Mahul Chakraborty (UC Irvine) for their unpublished chromosome level assembly of a Texas individual of *H. c. vasquezae.* Benita Laird-Hopkins (University of South Bohemia) for providing *H. charithonia* samples from Panama. Elena Mistreanu for her assistance with greenhouse work. The authors also thank the Butterfly Ecology and Evolution Research group, in particular Carlos Arias, Rémi Maxion, Jessie Foley and Laura Hebberecht for all their support during our stay at STRI (Gamboa, Panama).

## Author contributions

F.S. and J.M conceived this study. F.S. performed the analysis of genomic data; F.S. and S.D. performed laboratory work; F.S. and S.G. designed and performed the cross experiments; F.S. and J.M. wrote the manuscript with contributions from the other authors.

