## Supplementary Figures for "Ancient inversion polymorphisms are locally adaptive in a widespread butterfly species"

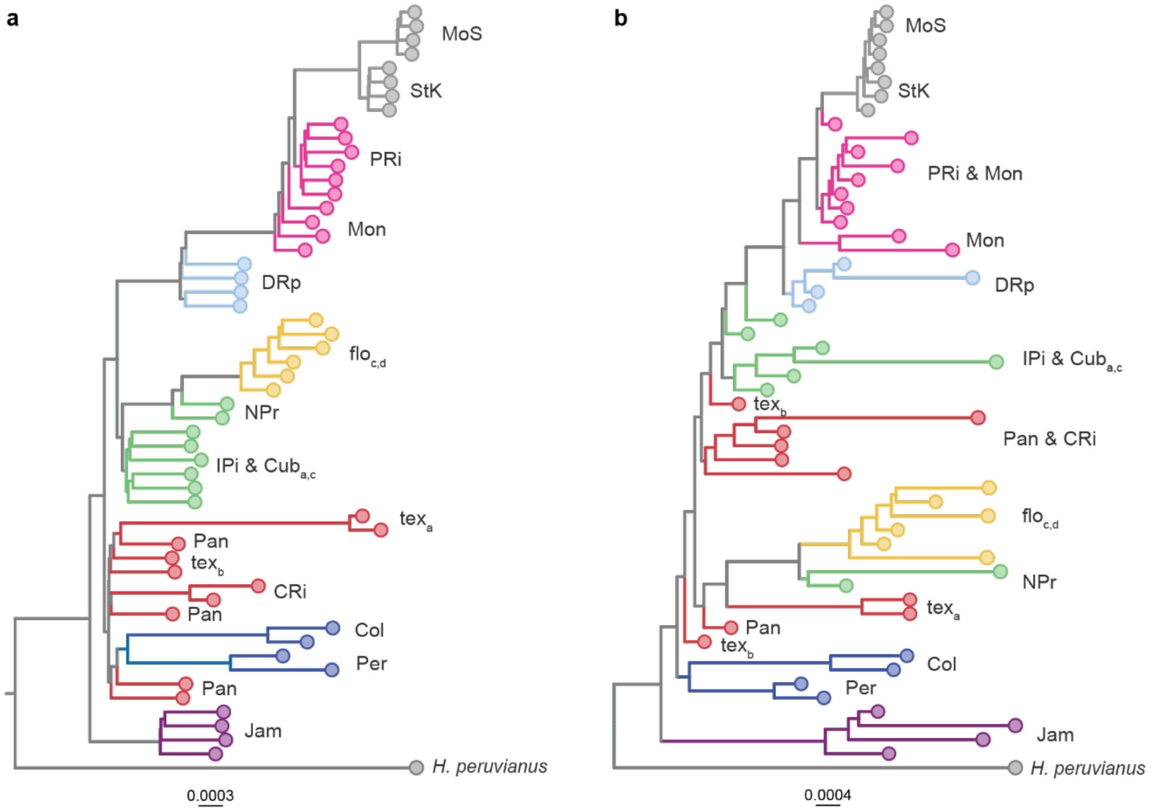


**Supplementary Figure 1** – Neighbor-Joining (NJ) tree based on **(a)** 269,403 autosomal sites sampled every 1-kb and **(b)** 12,886 Z-chromosome sites sampled every 1-kb.

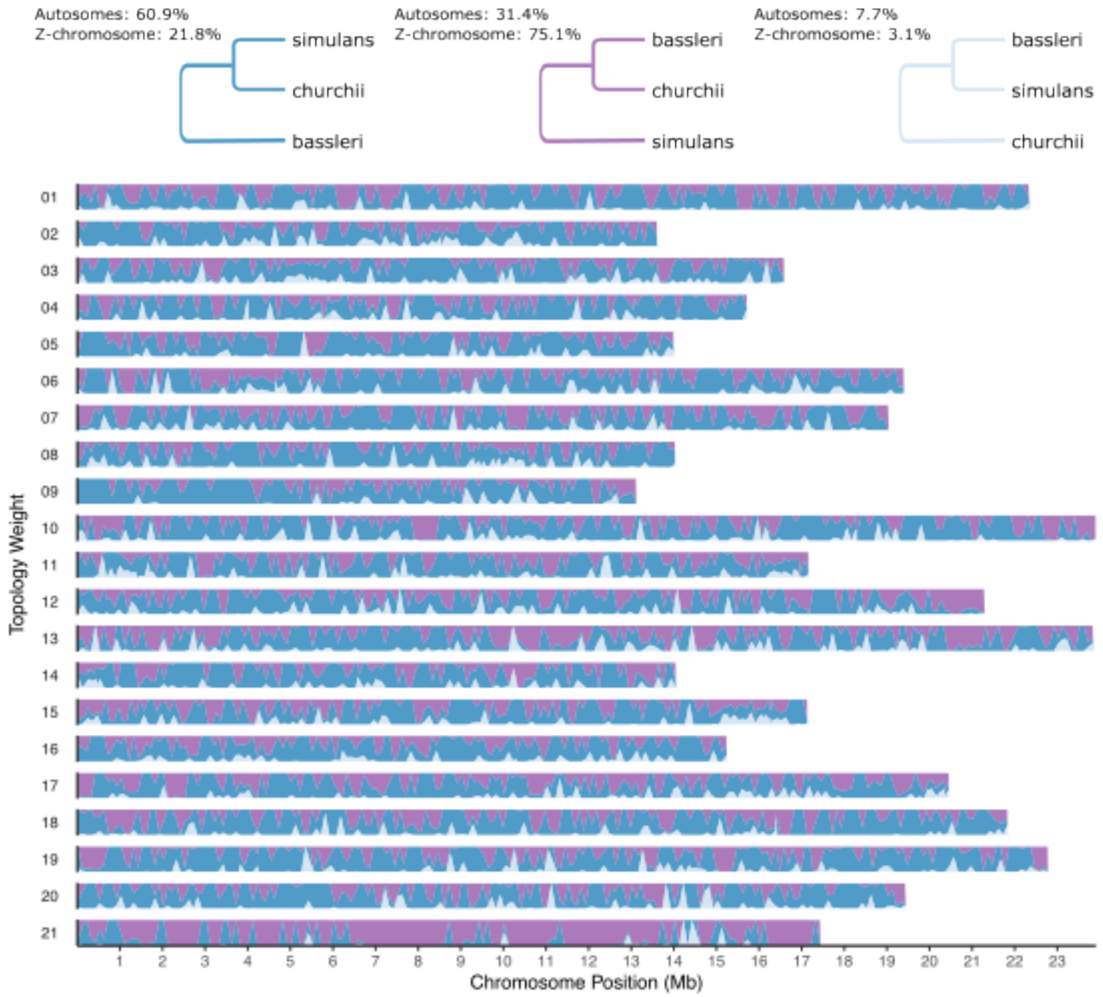


**Supplementary Figure 2 –Subspecies relationships along chromosomes.** Topology weightings were estimated using Twisst, in non-overlapping 50 kb window and smoothed as a locally weighted average. The three possible topologies and their respective average weights across autosomal and Z-chromosome windows, are indicated on top.

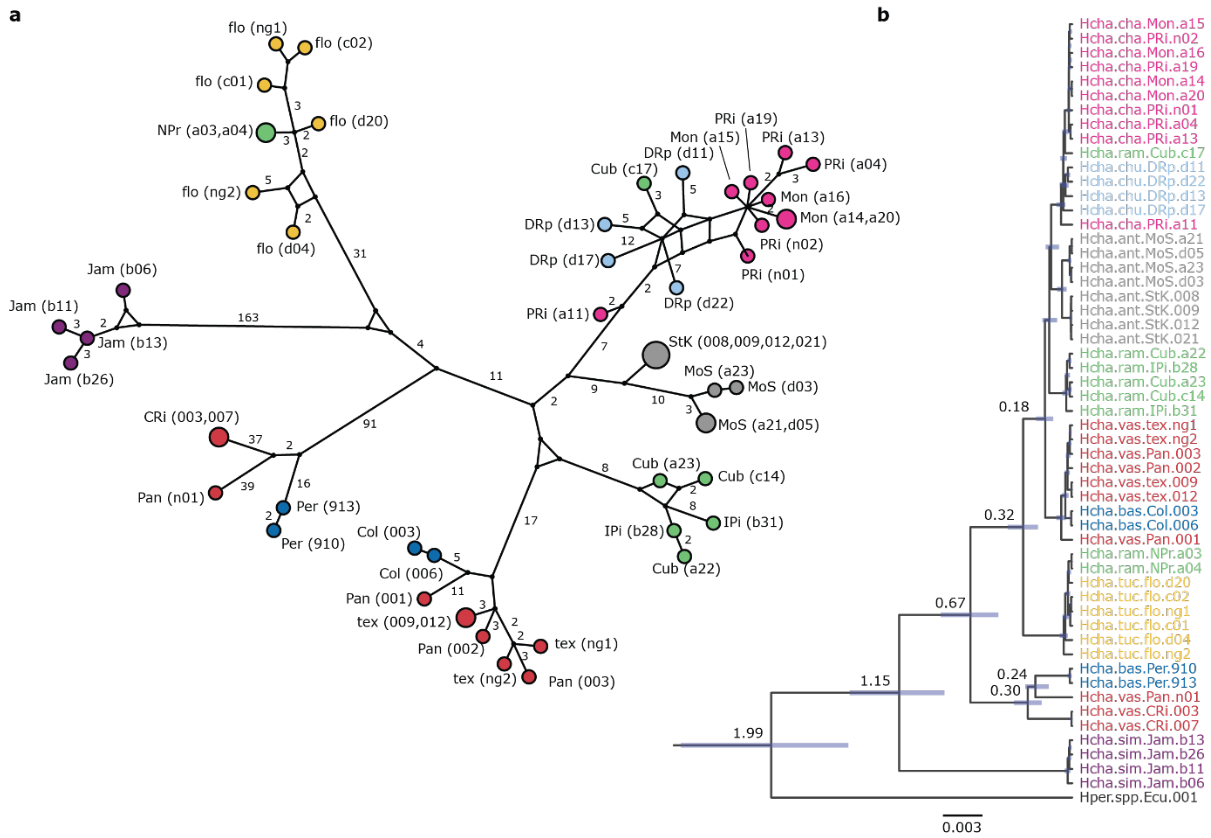


**Supplementary Figure 3 – Mitochondrial haplotype network and Bayesian phylogenetic tree suggest at least two colonization waves of the Caribbean islands. (a)** Median-joining network of concatenated mitochondrial gene alignments. Circle sizes are proportional to the number of individuals with that same haplotype and are colored according to subspecies. Location codes and individuals' codes within these locations are provided next to haplotypes. Mutational steps along edges are shown except for edges with only a single mutation. **(b)** Mitochondrial Bayesian tree estimated in BEAST. Node ages are depicted in million years (Mya) and were calibrated assuming a substitution rate of $1.15 \times 10^{-8}$ substitutions/site/year of the Cytochrome c oxidase subunit 1 (COI) region [1]. Node bars indicate the 95% HPD intervals. Individuals are colored according to subspecies and their respective codes are provided in Supplementary Table 1.

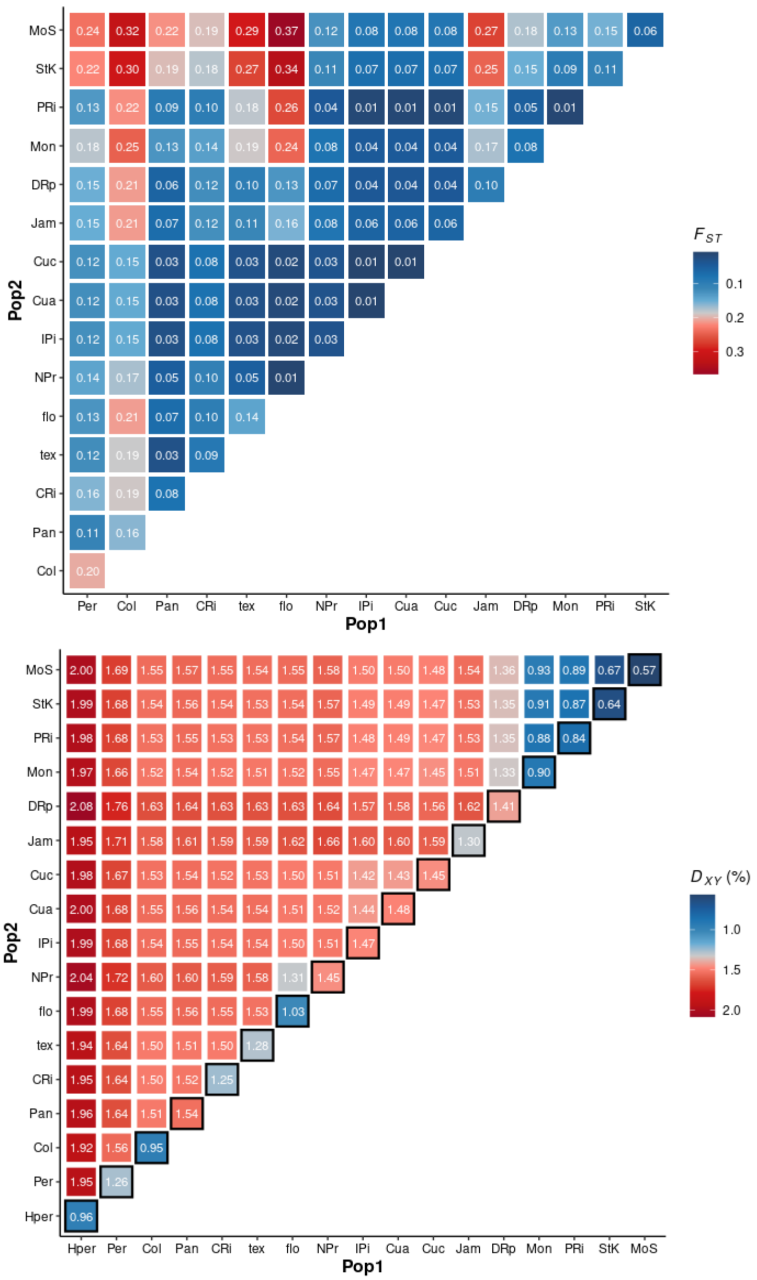


**Supplementary Figure 4 – Genome-wide levels of population divergence and diversity. (a)** Mean genome-wide $F_{ST}$, calculated in 50 kb non-overlapping windows. **(b)** Mean genome-wide nucleotide diversity (π; diagonal) and pairwise population absolute divergence ($d_{xy}$; off-diagonal), calculated in 50 kb non-overlapping windows. Population codes are provided in Supplementary Table 1.

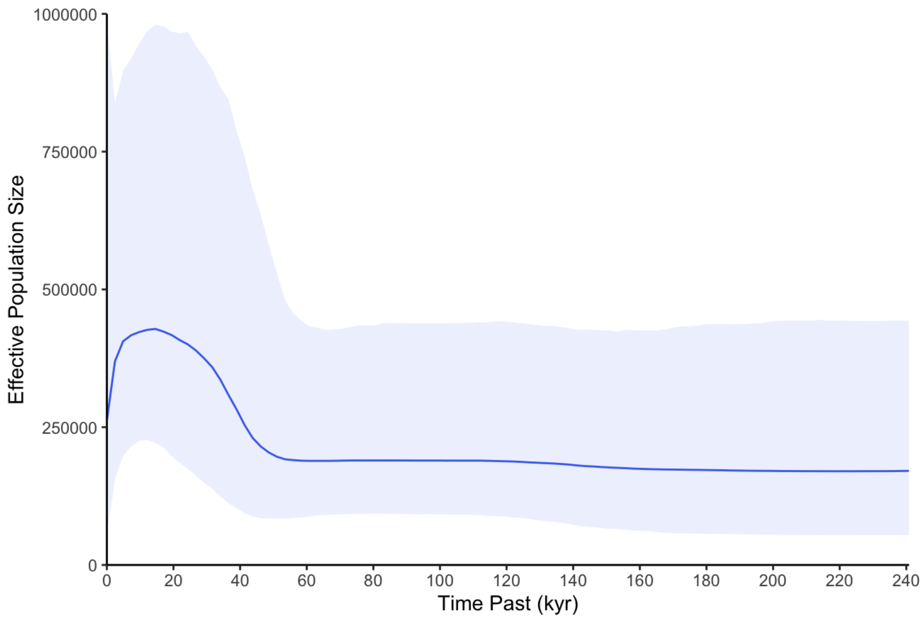


**Supplementary Figure 5** - Bayesian skyline plot (BSP) of *H. charithonia* mitochondrial haplotypes (excluding individuals from Jamaica). The solid blue line represents the median effective population size and the blue band the 95% high posterior density (HPD) interval. On the y-axis, the effective population sizes ($N_e$) are scaled to the mutation rate, whereas the x-axis represents coalescent time in thousands of years (kyr) (mutation rate and coalescent time based on ?Brower 1994 again?).

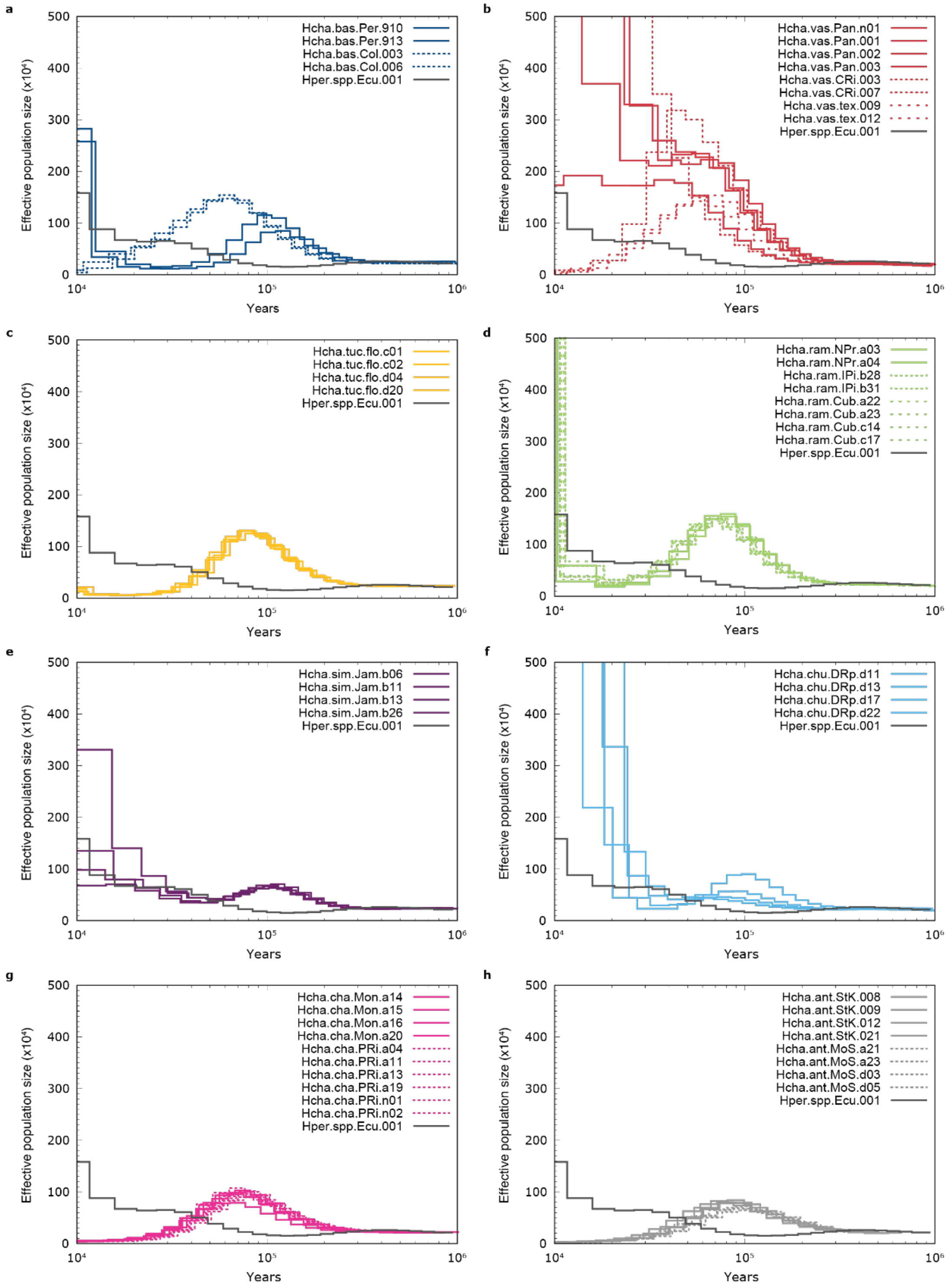


**Supplementary Figure 6** – Reconstruction of the past demographic history of *H. charithonia* using the Pairwise Sequentially Markovian Coalescent (PSMC) model from autosomal data. Times were calibrated assuming a substitution rate of $2.9\times10^{-9}$ substitutions/site/generation and a generation time of 0.25 years [2].

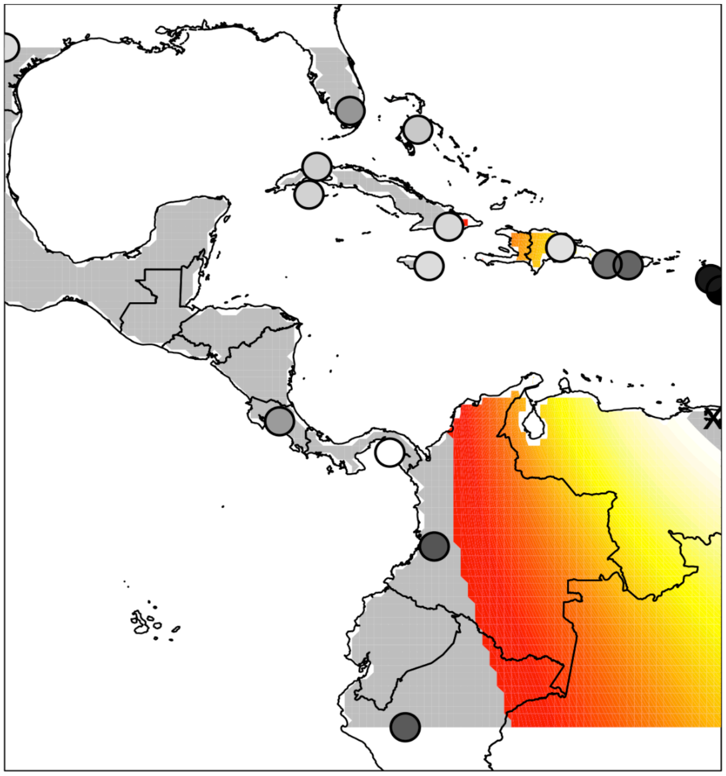

**Supplementary Figure 7 –Range expansion**. The directionality index (ψ) detects a range expansion with origin in Venezuela (X mark). Brighter yellow regions indicate more likely origin of expansion, while red and gray regions indicate unlikely origin. Each dot represents a sampled population and fill color indicate levels of diversity – darker grey indicates lower diversity.

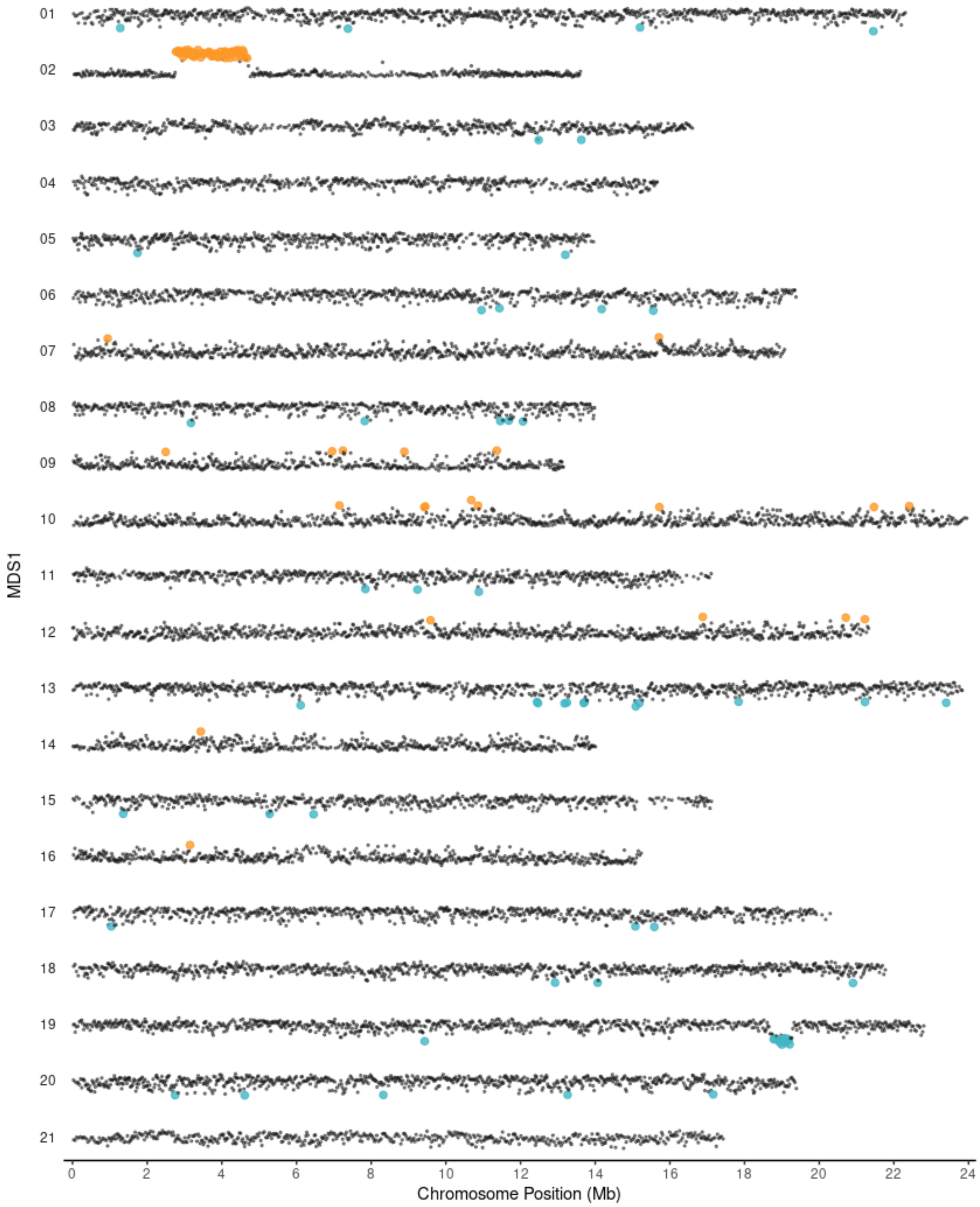


**Supplementary Figure 8 – Local PCA MDS plots along chromosomes.** Each dot represents a 500 SNP window (median block size = 22,852 bp), and windows with outlier MDS scores as measured by z-score are highlighted in orange (z-score > 3) and blue (z-score < -3).

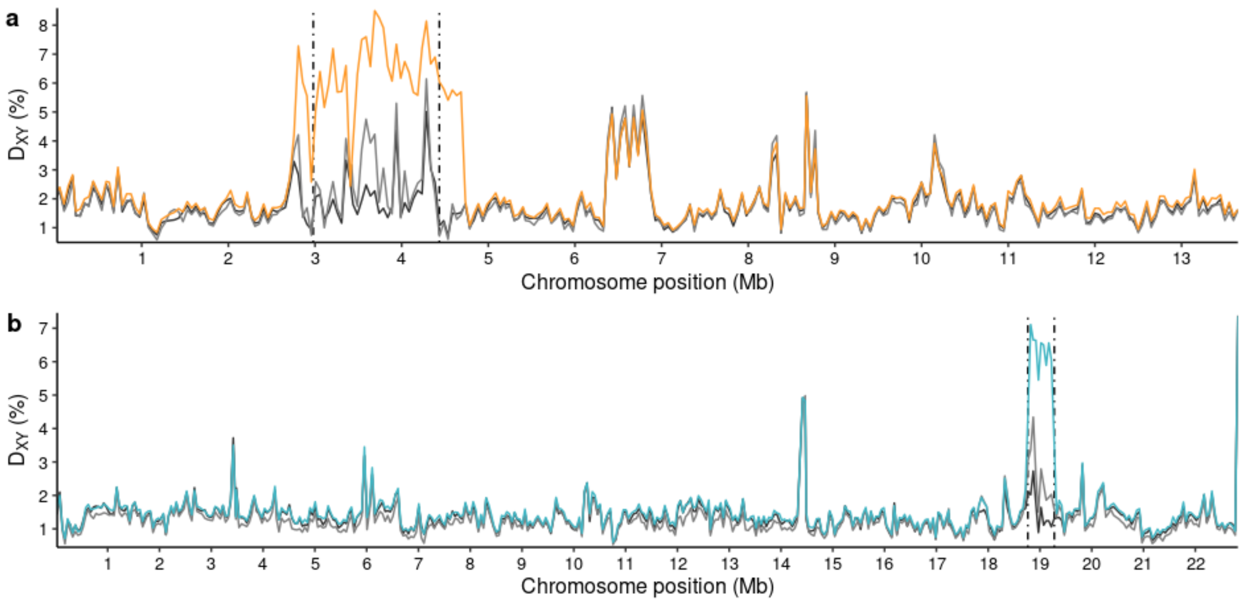


**Supplementary Figure 9 - Absolute divergence ($d_{xy}$) along chromosomes 2 (a) and 19 (b).** $d_{xy}$ was calculated in non-overlapping 50 kb windows, between individuals homozygous for different haplotypes at the putative inversions (orange, blue) and between individuals homozygous for the same haplotype (i.e., homozygous for the most frequent (dark grey) and least frequent (light grey) putative inversion haplotype). Individuals' genotypes were defined following the PCA of the putative inversions regions (Figure 2c,d). The dashed-dotted lines indicate the putative inversion breakpoints according to the local PCA (Supplementary Figure 8; Figure 2a,b).

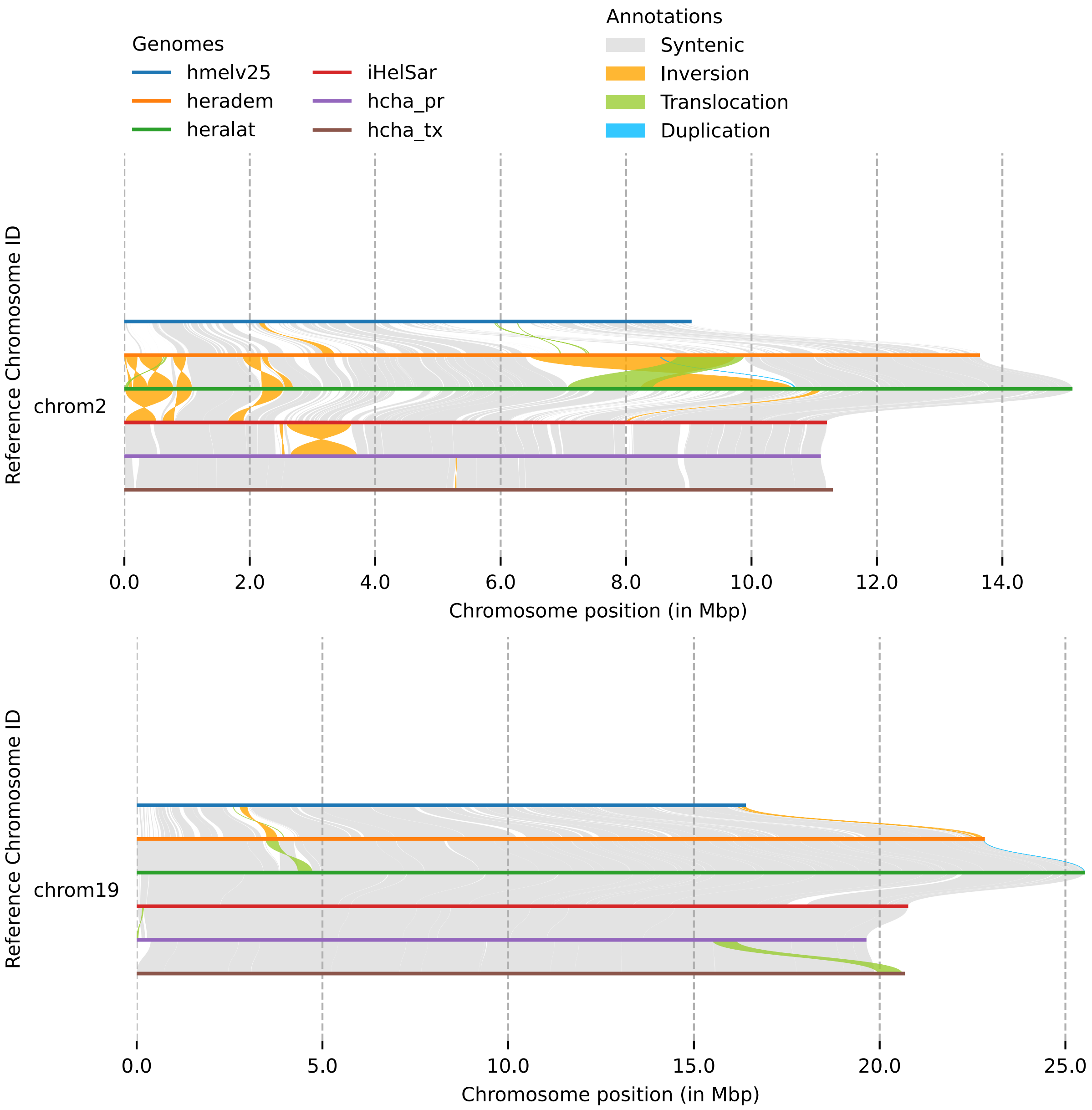


**Supplementary Figure 10 – Detection of structural rearrangements from chromosome alignments.** Species codes: *H. melpomene* (hmelv25); *H. erato demophoon* (heradem); *H. erato lativitta* (heralat); *H. sara* (iHelSar); *H. charithonia* Puerto Rico (hcha_pr); *H. charithonia* Texas (hcha_tx)

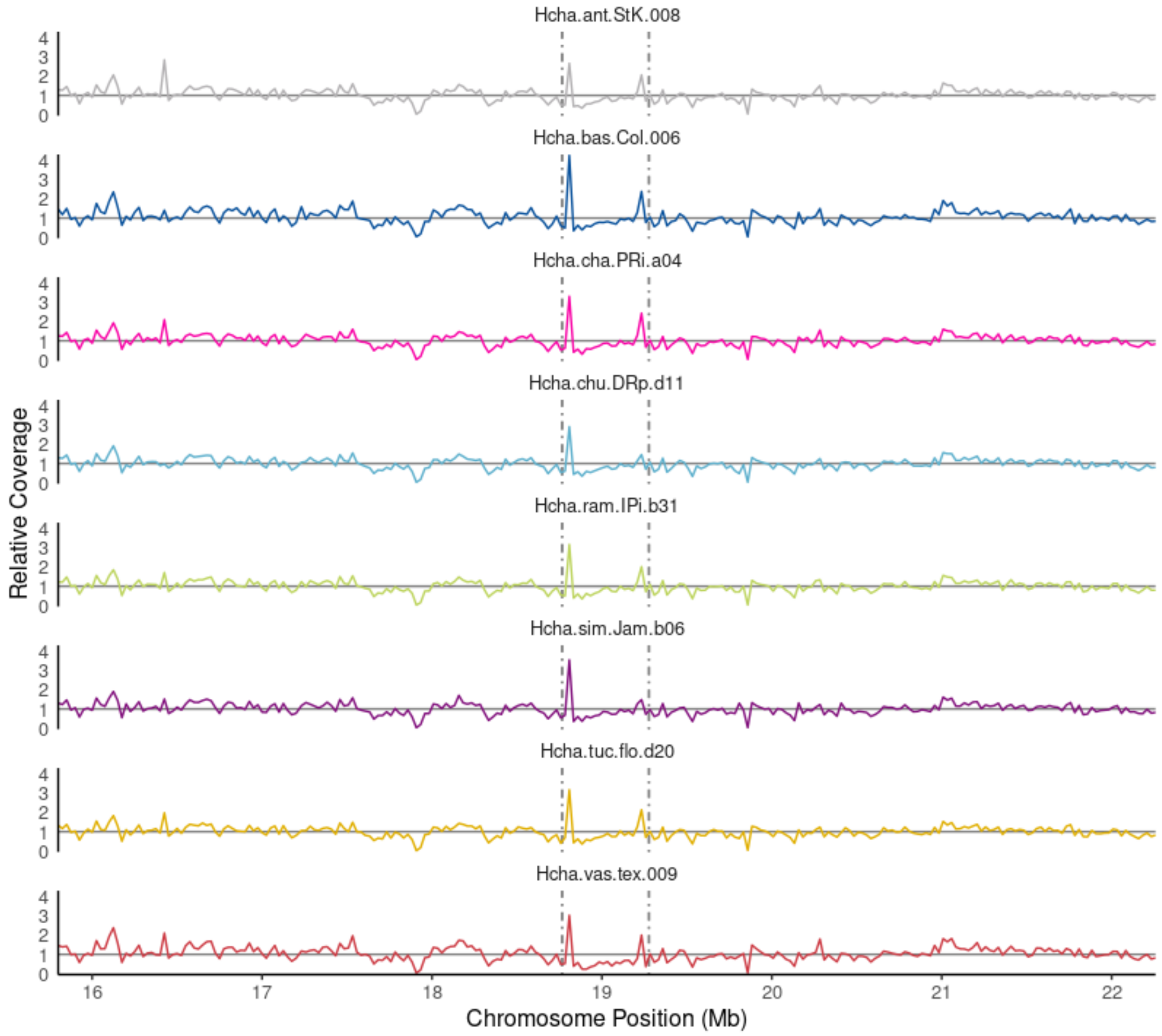


**Supplementary Figure 11 – Relative coverage along chromosomes.** Coverage along chromosome was estimated in sliding windows of 50 kb, and relative coverage was obtained by dividing each window coverage by the median coverage across all autosomal windows. The dashed-dotted lines indicate the putative inversion breakpoints according to the local PCA (Supplementary Figure 8; Figure 2a,b). Individuals are colored according to their respective subspecies assignment (see Figure 1; Supplementary Table1).

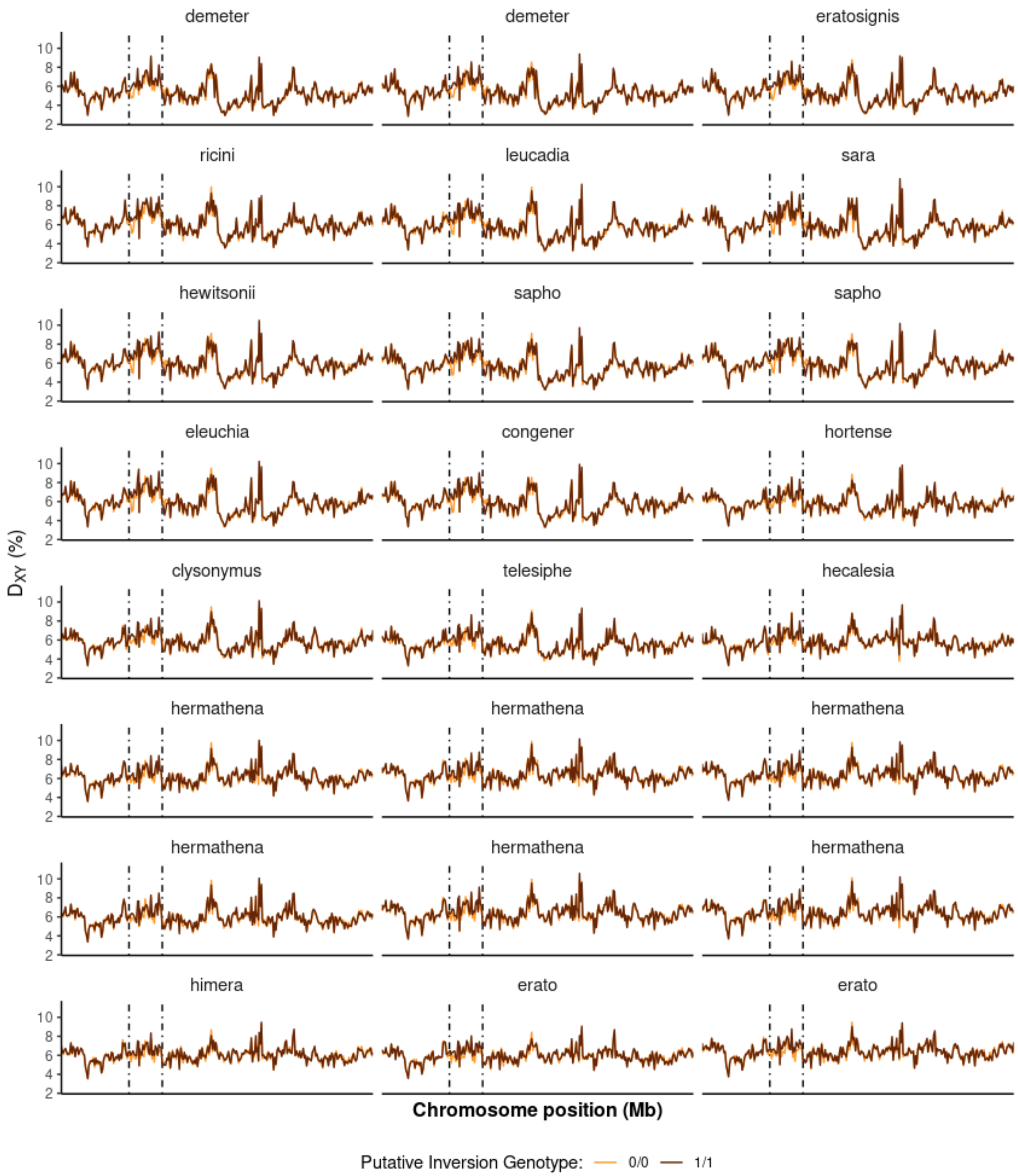


**Supplementary Figure 12 - Absolute genetic distance ($d_{xy}$) to outgroup species along chromosome 2 (50 kb non-overlapping windows)**. $d_{xy}$ was calculated between outgroup species and either one *H. charithonia* individual homozygous for the inversion (dark brown) or one *H. charithonia* individual homozygous for the standard haplotype (orange). The dashed-dotted lines indicate the putative inversion breakpoints according to the local PCA (Supplementary Figure 8; Figure 2a,b).

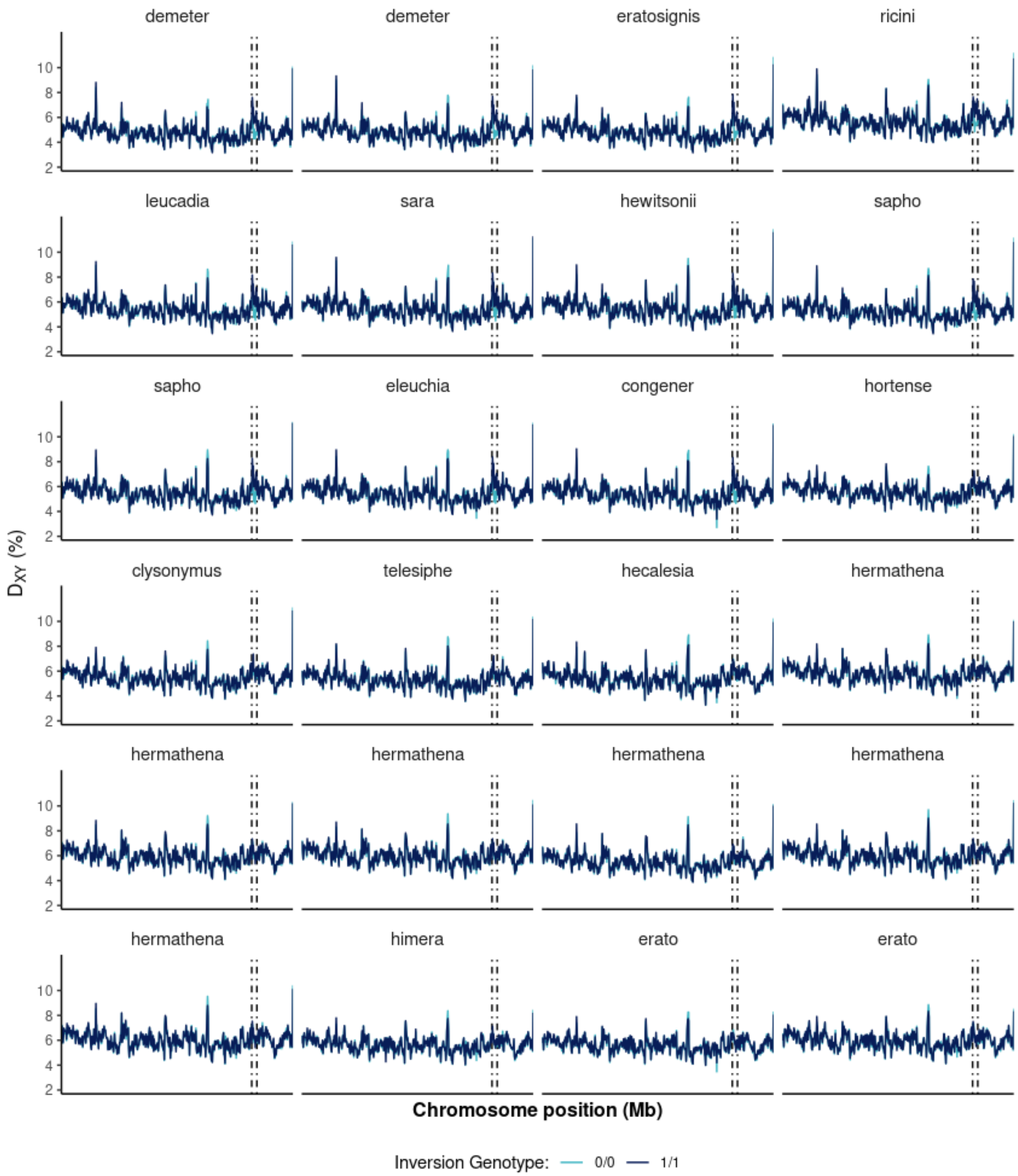


**Supplementary Figure 13 – Absolute genetic distance ($d_{xy}$) to outgroup species along chromosome 18 (50 kb non-overlapping windows)**. $d_{xy}$ was calculated between outgroup species and either one *H. charithonia* individual homozygous for the inversion (dark blue) or one *H. charithonia* individual homozygous for the standard haplotype (light blue). The dashed-dotted lines indicate the putative inversion breakpoints according to the local PCA (Supplementary Figure 8; Figure 2a,b).

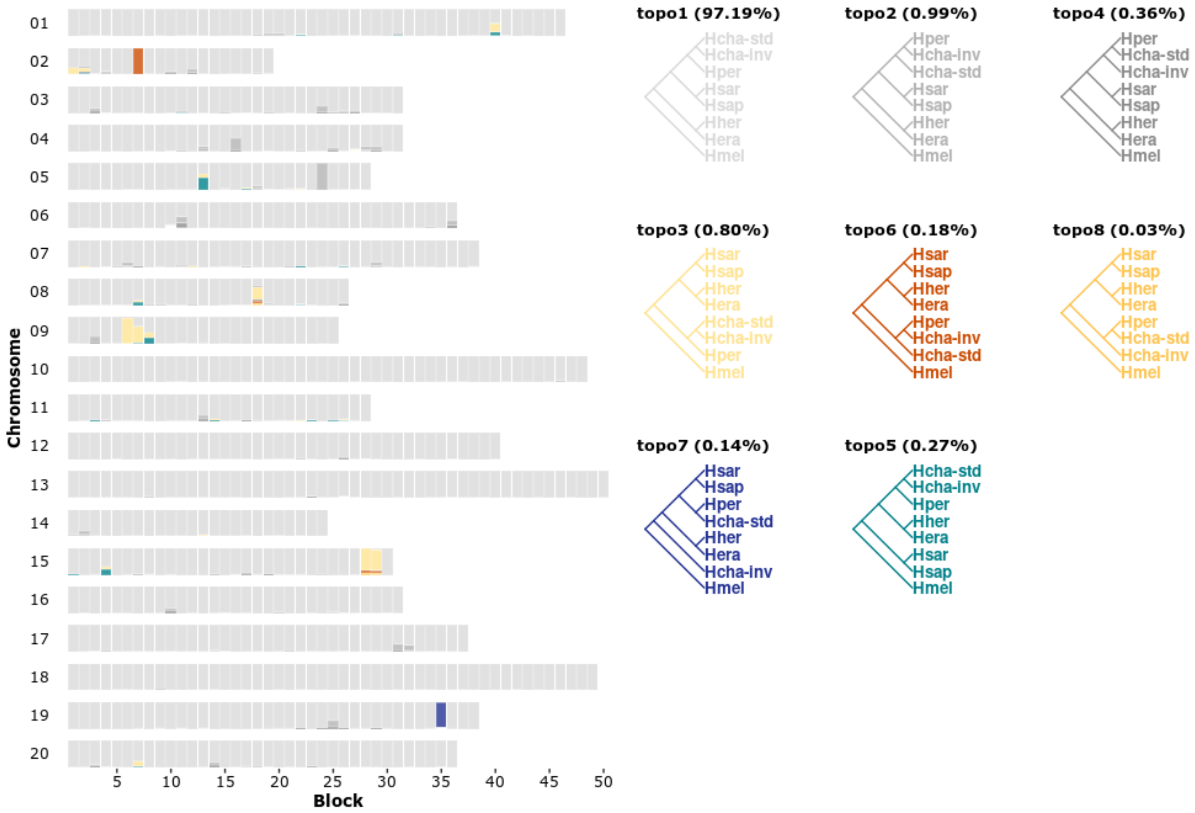


**Supplementary Figure 14 – Estimated species trees across chromosomes under the multispecies coalescent (BPP analyses A01).** Each bar represents one block of 100 loci. For each block the posterior probabilities of different trees are given in different colors. Only the eight most frequent trees are shown, and posterior probabilities of the remaining trees are shown in white.

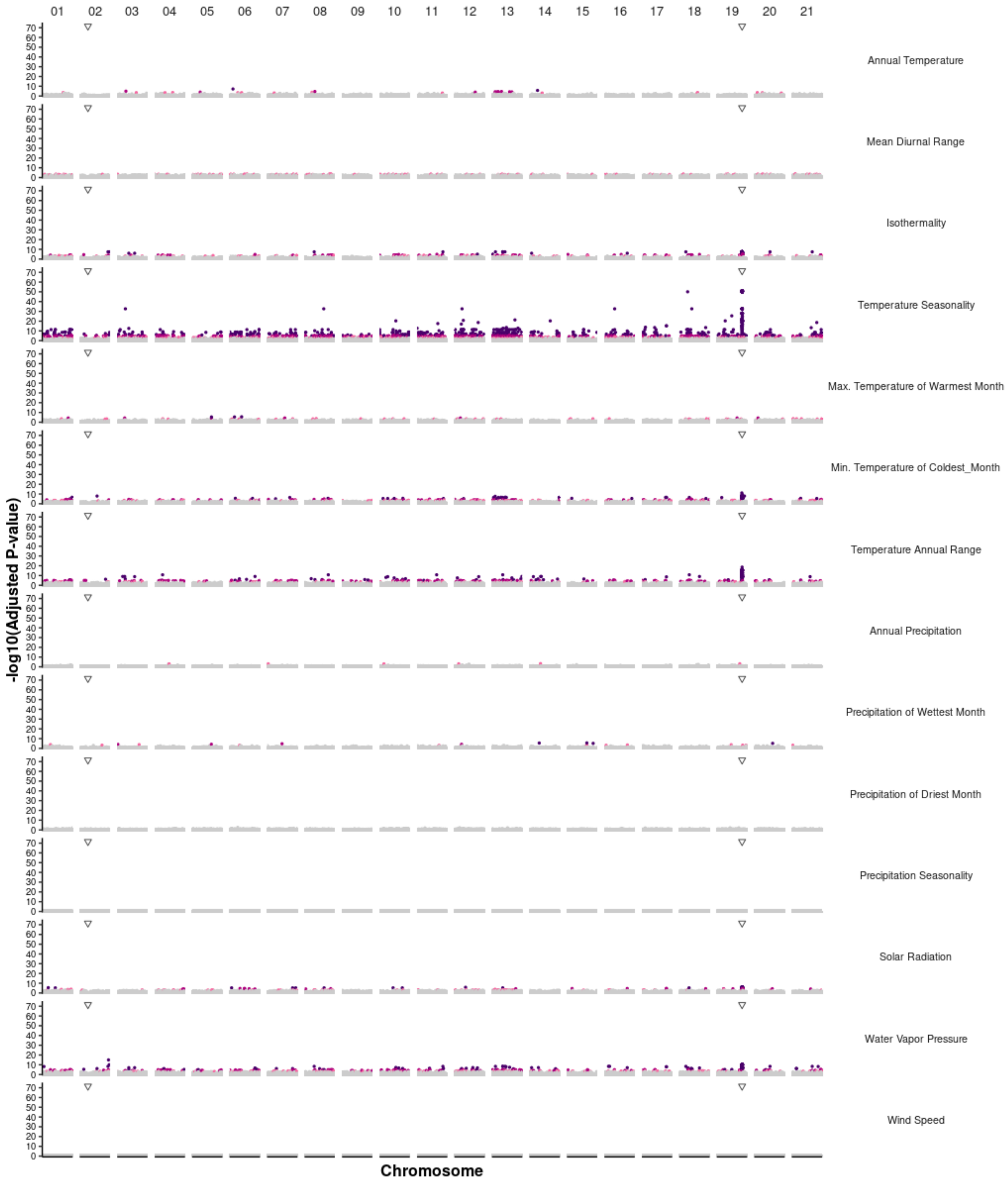


**Supplementary Figure 15 – Genotype-environment associations.** The Manhattan plot shows the P values from the latent factor mixed models (LFMM). Points are colored according to FDR (dark brown: <0.00001, dark orange: <0.0001, orange: <0.001). Inverted triangles represent the inferred location of inversions.

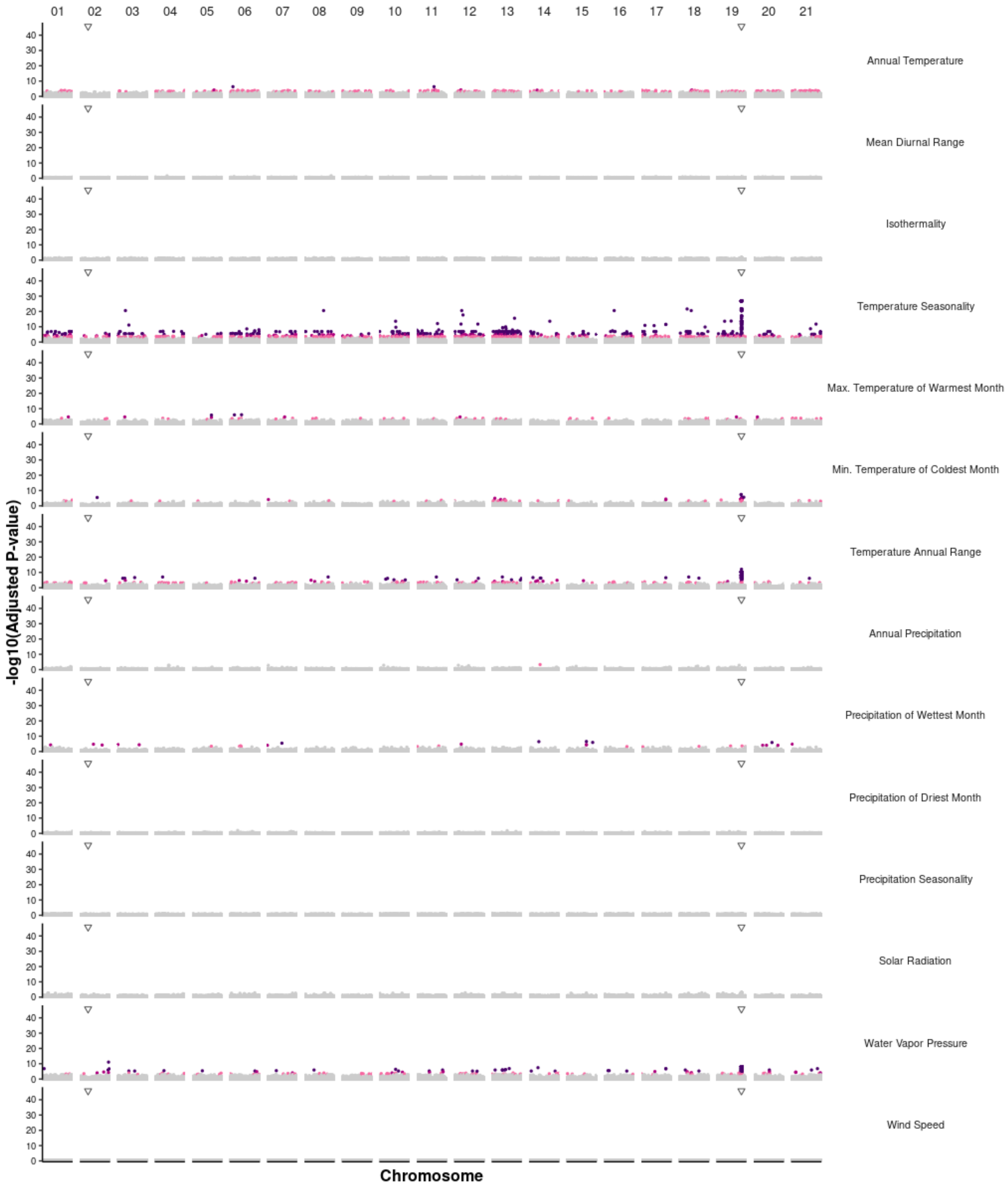


**Supplementary Figure 16 - Genotype-environment associations.** Given each summer Texas is colonized by *H. charithonia* migrants from Mexico, we took a conservative approach and considered climatic values extracted from Monterey (Mexico; latitude: 25.7, longitude: -100.3) for the Texas' individuals. This location has all year-round resident populations of *H. c. vazquezae*. The Manhattan plots showing association with climatic variables along chromosomes. Each point indicates P-values at each SNP. Points are colored according to their FDR values (purple: <0.00001, dark pink: <0.0001, light pink: <0.001; grey: > 0.001). The dashed-dotted vertical lines represent the inferred location of inversion breakpoints.

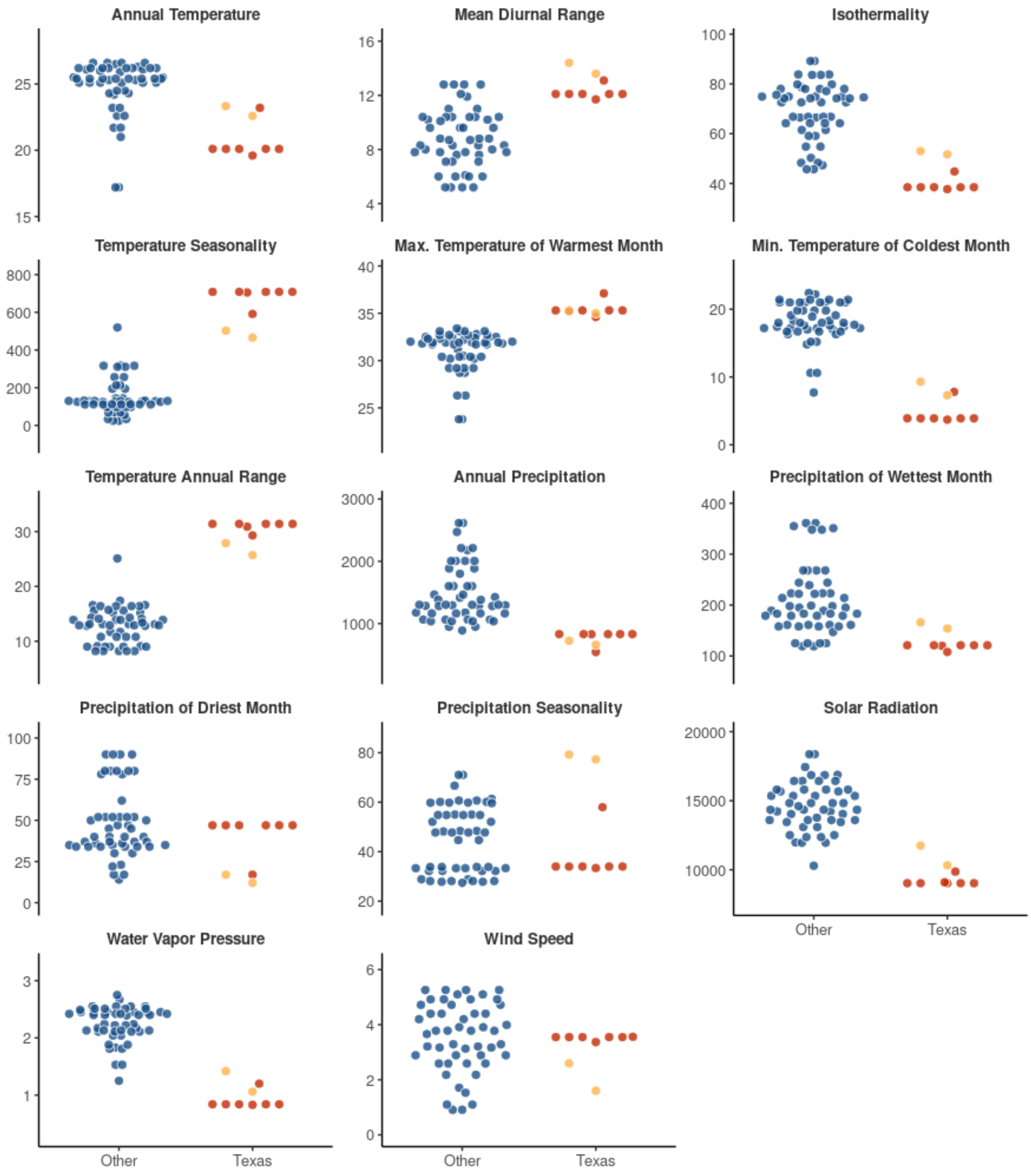


**Supplementary Figure 17 – Environmental conditions at locations with and without the chromosome 19 inversion genotype.** Each dot represents one individual. For each individual, the environmental values at its geographical location were recorded (y-axis). Individuals are plotted according to whether they were sampled in Texas, or elsewhere in *H. charithonia* distribution, which also corresponds to their genotype at chromosome 19 inversion: Texas (inverted genotype, red), Other (standard genotype, blue). The two yellow dots represent pseudo-locations south of Texas, with all year-round resident populations from which individuals migrate north during warmer periods: Tamaulipas (Mexico: latitude: 23.7, longitude: -99.1) and Monterey (Mexico; latitude: 25.7, longitude: -100.3).

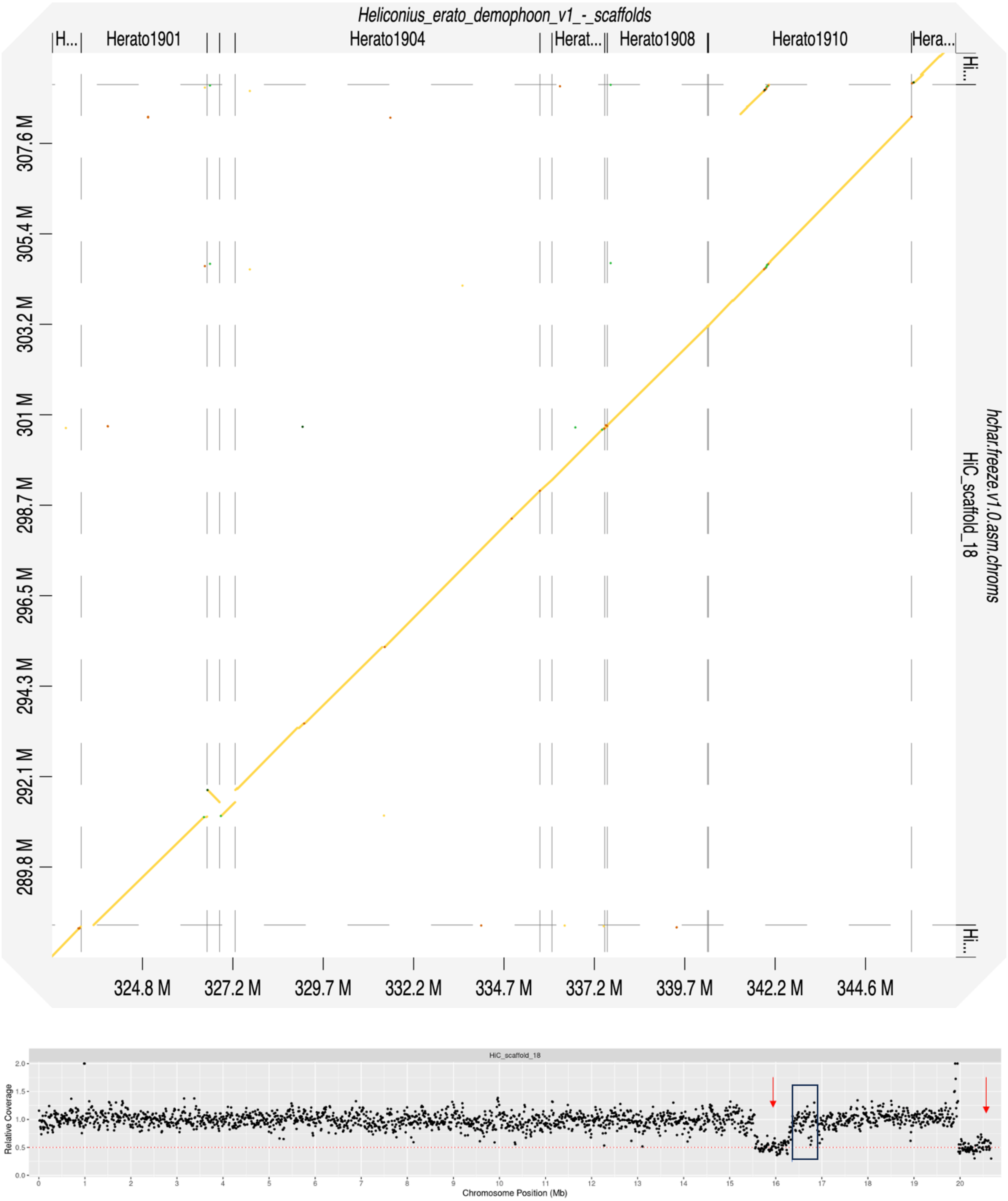


**Supplementary Figure 18 – Texas *H. charithonia* genome mis-assembly on chromosome 19.** D-genies genome-genome alignment (top panel) shows two genomic blocks in *H. charithonia* (Texas) genome assembly map to the same region in *H. erato*, immediately adjacent to the putative inversion. Relative coverage along chromosome 19 (bottom panel), shows the two blocks (indicated by arrows), have half of the expected coverage and likely correspond to divergent haplotypes that were assembled separately. The rectangle indicates the approximate coordinates of the putative inversion.
